# Oxidative and non-oxidative active turnover of genomic methylcytosine in distinct pluripotent states

**DOI:** 10.1101/846584

**Authors:** Fabio Spada, Sarah Schiffers, Angie Kirchner, Yingqian Zhang, Gautier Arista, Olesea Kosmatchev, Eva Korytiakova, René Rahimoff, Charlotte Ebert, Thomas Carell

## Abstract

Epigenetic plasticity underpins cell potency, but the extent to which active turnover of DNA methylation contributes to such plasticity is not known and the underlying pathways are poorly understood. Here we use metabolic labelling with stable isotopes and mass spectrometry to quantitatively address the global turnover of genomic methylcytidine (mdC), hydroxymethylcytidine (hmdC) and formylcytidine (fdC) across mouse pluripotent cell states. High rates of mdC/hmdC oxidation and fdC turnover characterize a formative-like pluripotent state. In primed pluripotent cells the global mdC turnover rate is about 3-6% faster than can be explained by passive dilution through DNA synthesis. While this active component is largely dependent on Tet-mediated mdC oxidation, we unveiled an additional mdC oxidation-independent turnover process based on DNA repair. This process accelerates upon acquisition of primed pluripotency and returns to low levels in lineage committed cells. Thus, in pluripotent cells active mdC turnover involves both mdC oxidation-dependent and -independent processes.

## Introduction

Methylation at position 5 of deoxycytidine (dC) is the most prominent post-replicative nucleobase modification in higher eukaryotic genomes and its functional relevance has been extensively investigated^1,2^. In mammals, genomic dC is methylated to 5-methyl-2’-deoxycytidine (mdC) by the DNA (cytosine-5)-methyltransferases DNMT1, DNMT3a and DNMT3b^3^, mostly, but not exclusively in the context of CpG dinucleotides^4^. These enzymes differentially contribute to establishing and maintaining genomic mdC patterns across cellular and organismal generations. In addition, genomic mdC can be further modified by iterative oxidation to 5-hydroxymethyl-, 5-formyl-and 5-carboxyl-dC (hmdC, fdC and cadC, respectively) through the action of Ten-Eleven Translocation (TET) di-oxygenases^5–7^. As several classes of DNA binding factors display differential affinity for their target sequences depending on the dC modification state^2,1,8^, genomic patterns of dC modification contribute to consolidation and propagation of gene expression states and chromatin structure, thus affecting fundamental processes such as cell fate determination and restriction, imprinting, X chromosome inactivation and genome stability^1,2^.

In the last two decades methylation of genomic dC was shown to be rapidly removed in various biological contexts^9^. Several erasure mechanisms were proposed, including prevention of methylation maintenance, which results in passive dilution of the modified base through DNA replication, and active demethylation by enzymatic processes^9^ (Fig. 1a). Among the latter oxidation of mdC to fdC or cadC by TET enzymes followed by base excision repair (BER) mediated by the DNA glycosylases TDG^9,10^ and Neil1/2^11,12^ is best characterized. Additional proposals invoke the involvement of other DNA repair pathways^9^ or direct C-C bond cleavage^13–16^. A third line of proposal posits that genomic mdC and hmdC could be deaminated to thymidine (dT) and 5-hydroxymethyl-2’-deoxyuridine (hmdU), respectively, either by a cytosine deaminase of the AID/APOBEC family^9^ or by *de novo* DNA methyltransferases DNMT3a/b^17^. The resulting T:G or hmU:G mismatches would be then resolved by BER or non-canonical mismatch repair (ncMMR)^18–21^. Finally, it was suggested that unmodified dC could be deaminated by AID and that long patch BER or ncMMR could lead to indirect co-removal of adjacent ^m^C residues^20–22^. Currently, all these pathways are highly controversial, as there is no conclusive evidence for their functional relevance *in vivo*^9^. It is however conceivable that different pathways operate at different stages during development.

**Figure 1.**
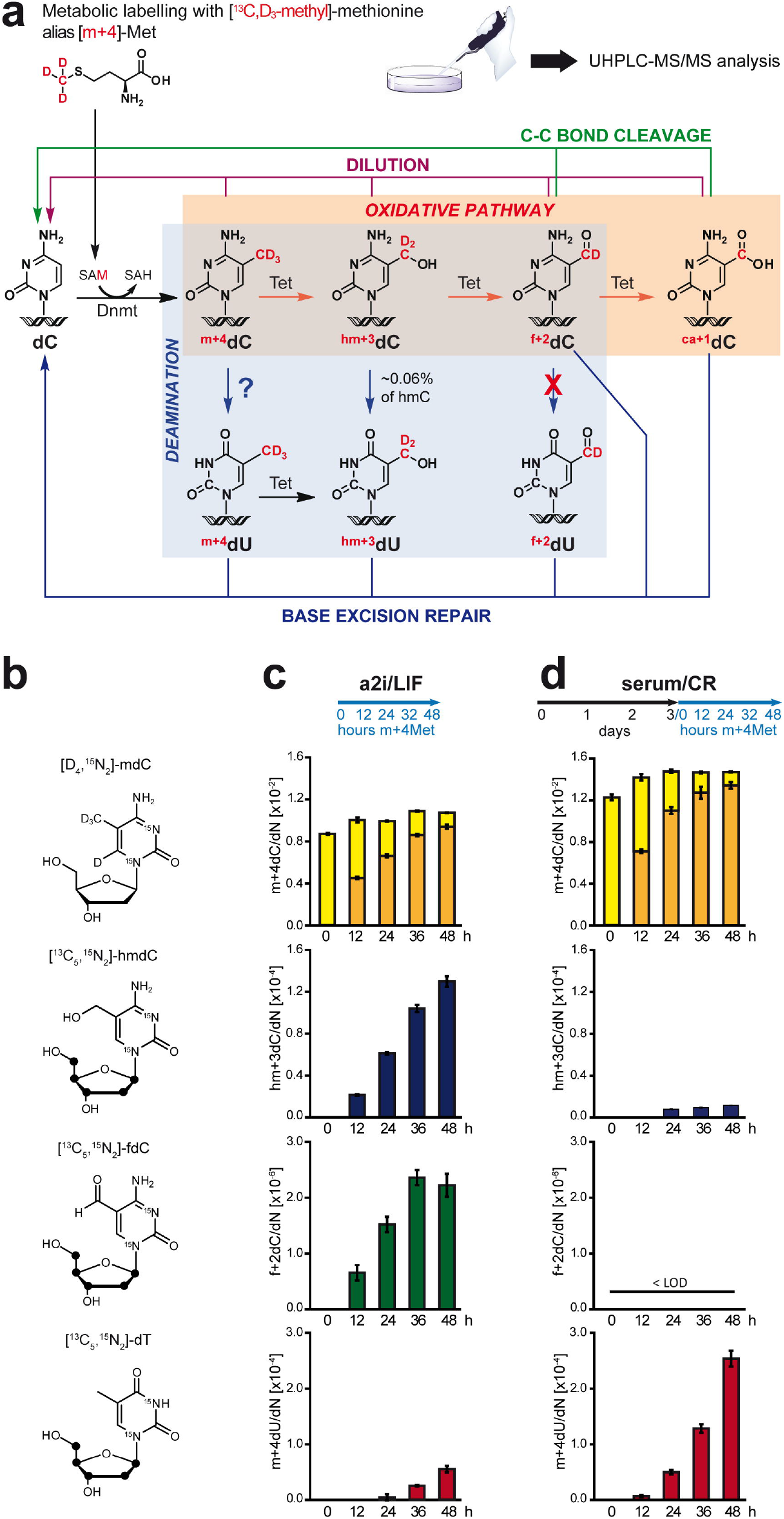
Drastic reduction of global genomic mdC oxidation and increase of mdC to mdU conversion upon transition to primed pluripotency. **a**) Proposed de-modification pathways for genomic C5-cytosine modifications and expected labelled mdC derivatives upon metabolic labelling with m+4Met. **b**) Nucleoside isotopologues used as internal standards for UHPLC-MS^2^ analysis. Black balls in the ribose represent ^13^C atoms. **c** and **d**) Time course analysis of genomic dC derivatives upon metabolic labelling with m+4Met under naïve (c; a2i/LIF) and priming (d; serum/CR) conditions. The labelling/time course schedules are shown at the top. Global levels of unlabelled (pale yellow) mdC and m+4dC (dark yellow), hm+3C (blue) and f+2C (green) are shown as mean and standard deviation of three technical replicates. See also Fig. S3 for an independent biological replicate. LOD = limt of detection.

During early embryonic development, the epiblast transits through a spectrum of pluripotent states^23^. This transition can be modelled by cultivating mouse pluripotent stem cells (mPSCs) under specific conditions and involves dynamic changes in expression and epigenetic landscapes, including a major global gain of mdC^23^. Conversely, reversion from partially and heterogeneously primed mPSC cultures to the naïve state involves downregulation of *de novo* methyltransferases and collapse of DNA methylation maintenance with consequent global and passive (Tet enzyme-independent) loss of genomic methylation^24^. In contrast, little is known about the extent and nature of the turnover of modified genomic cytosines during the forward, developmentally relevant transition from naïve to primed pluripotent states. Here, we used metabolic labelling with stable isotope standards for highly accurate mass spectrometry based quantification of the global genomic mdC, hmdC and fdC turnover through a progression of pluripotent states. We show that upon acquisition of primed pluripotency roughly 3-6% of the methylome is actively turned over. Most of this active turnover is due to Tet-mediated mdC oxidation, while a minor part involves an unknown mdC oxidation-independent DNA repair process.

## Results

### Global genomic mdC oxidation kinetics decrease upon progression to primed pluripotency

We cultured mPSCs under specific conditions to model the progression of the embryonic epiblast from naïve to primed pluripotency and investigate the kinetics and turnover of dC modification during this transition. In order to monitor priming conditions, we first made use of a mPSC line expressing an Oct4-YFP reporter^25^. After adaptation to naïve conditions in the presence of Gsk3 inhibitor CHIR99021 (CHIR), Mek1/2 inhibitor PD0325901 (also known as 2i) and LIF, the cells were plated at low density in serum-containing medium supplemented with CHIR and the Tankyrase inhibitor IWR1-endo (serum/CR) as previously described^26^. Live cell microscopy over the course of 6 days without passaging showed that these cells rapidly formed monolayer colonies that retained a remarkably homogeneous nuclear YFP signal across the culture (Supplementary Fig. S1a), indicating that under these conditions mPSC cultures may not loose pluripotency. Similarly intense and homogeneous YFP signals were observed when the same reporter line was permanently primed to mouse Epiblast Stem Cells (mEpiSCs) in serum-free, chemically defined medium supplemented with CHIR, IWR1, FGF-2 and Activin A (CDM/CRFA; Supplementary Fig. S1a). Expression analysis of pluripotency factors in serum/CR cultures and CDM/CRFA EpiSCs revealed similarly reduced and increased transcript levels of naïve and primed pluripotency factors, respectively (Supplementary Fig. S1b), showing that these conditions support similarly primed states. Thus, even without prolonged passaging or in chemically defined conditions, CHIR/IWR1 treatment supports primed pluripotency. We then used ultra-high pressure liquid chromatography coupled with triple quadrupole mass spectrometry (UHPLC-MS^2^)^27^ to analyse global levels of genomic mdC and hmdC upon priming of mPSCs under the same conditions over a five day time course. As previously reported^28^, the transition from the naïve (2i/LIF) to a homogeneously primed pluripotent state was marked by a large increase in global levels of genomic mdC, while hmdC abundance dropped sharply, both modifications reaching plateau levels by day 4 (Supplementary Fig. S2a). Global levels of genomic mdC and hmdC similar to those in serum/CR cultures were detected in Oct4-YFP mEpiSCs under serum-free conditions, excluding potential confounding effects of undefined serum composition. Note that under both serum-containing and serum-free conditions we avoided supplementation with ascorbic acid, which was shown to greatly increase the levels of oxidized mdC derivatives^29^. The decrease in hmdC abundance upon priming is consistent with a recent proteomic analysis showing that the levels of chromatin bound Tet1 and 2 proteins are markedly reduced in mEpiSCs compared to naïve mPSCs, while Tet3 protein is undetectable in both states^30^.

Notably, under 2i/LIF conditions, mPSCs exhibit low levels of genomic cytosine modification^23^, while replacement of the Mek inhibitor with the Src inhibitor CGP77675 (conditions referred to as alternative 2i or a2i) results in relatively high genomic mdC levels^31,32^. Nevertheless, gene expression patterns remain similar and developmental potential unaltered relative to those of naïve mPSCs^31,32^. In particular, our expression analysis showed that the pattern of pluripotency and early lineage specification factors in mPSCs under a2i/LIF conditions was intermediate to that of naïve and primed states, with still high levels of Tfcp2l1 and Klf4 transcripts relative to 2i/LIF (naïve) conditions, but intermediate expression of Rex1, Stella and Prdm14 as well as low levels of Fgf5, Oct6, Otx2 and Dnmt3b relative to primed conditions (Fig. S1b). This pattern is highly similar to that previously shown for formative pluripotency, an intermediate state along the trajectory from naïve to primed pluripotency, during which competence for multi-lineage commitment is acquired^33^. In addition, high depth bisulfite amplicon sequencing revealed increased *de novo* methylation at secondary imprints in the transition from a2i/LIF to serum/CR conditions (Supplementary Fig. S2b), which is similar to the progression previously shown from pre-to early post-implantation stages in mouse embryos^34^. Altogether, our expression and genomic methylation analyses support that cultures transited from a2i/LIF to serum/CR conditions recapitulate fundamental molecular processes of the progression to a genuine primed pluripotent state. Importantly, prolonged culture under 2i/LIF conditions leads to loss of imprinted methylation and genetic instability, both are persevered in a2i/LIF long term cultures^31,32^. In addition, a2i/LIF and serum/CR conditions allow comparison of genomic cytosine modification dynamics between functionally distinct pluripotent states with substantial and comparable steady state levels of genomic mdC. Thus, we adopted a2i/LIF conditions both for routine mPSC maintenance and as starting state for further time course analyses of mPSC priming.

We then set out to monitor global cytosine modification kinetics during acquisition of primed pluripotency by metabolic labelling of the C5 substituent. To this aim we supplemented L-Methionine (Met)-free medium with isotopically labelled Met (m+4Met), whereby the methyl group provides a mass shift of four units ([*methyl*-^13^C,d3], hereafter referred to as m+4). As for naturally occurring Met, m+4Met is incorporated into S-adenosylmethionine (SAM) and m+4 is transferred thereof onto genomic dC by Dnmts, generating m+4dC. The latter can be sequentially oxidised by Tet proteins to hm+3dC, f+2dC and ca+1dC (Fig. 1a)^27^.

In order to attain accurate absolute quantification of both naturally occurring (mdC, hmdC, fdC) and heavy (m+4dC, hm+3dC, f+2dC) isotopologues, we synthesised an even heavier series of corresponding isotopologues ([d_4_,^15^N_2_]-mdC, [^13^C_5_,^15^N_2_]-hmdC and [^13^C_5_,^15^N_2_]-fdC; Fig. 1b) and used them as internal standards for quantification by UHPLC-MS^2^. In this work cadC and ca+1dC were not considered as isotopologues differing by only one mass unit cannot be reliably discriminated using UHPLC-MS^2^. We first compared the modification kinetics in cultures maintained constantly under a2i/LIF conditions to cultures undergoing priming in CHIR/IWR1 medium. In both cases we performed time course analyses over 2 days of labelling in the presence of m+4Met, which in the case of priming cultures was from day 3 to day 5 after transfer to serum/CR medium (Fig.s 1c,d, Supplementary Fig. S3). Relative to a2i/LIF conditions, in priming cultures the rate of m+4dC oxidation to hm+3dC was clearly slower. Accumulation of the further oxidized state f+2dC became undetectable under these conditions. Thus, the transition to primed pluripotency determines a major decrease in global oxidation of genomic mdC, which is consistent with the drastically reduced steady state levels of genomic hmdC (Supplementary Fig. S2a) and with a recent report of lower levels of chromatin bound Tet proteins in primed cultures^30^

### An active and largely Tet enzyme-dependent component of genomic mdC turnover in primed pluripotent cells

To monitor the turnover of modified genomic cytosines we reversed the labelling schema described above. We first pulsed labelled cultures with m+4Met, then chased the label for two days by transferring the cultures to medium with naturally occurring Met and collected samples every 12 h during the chase (Fig. 2a). We again compared cultures maintained constantly under serum/a2i/LIF conditions to cultures undergoing priming in serum/CR medium. In both cases the cultures were pulsed for five days with m+4Met before chasing. In the case of the primed cultures the first two pulse days were in a2i/LIF medium, while the last three were in serum/CR medium, thus corresponding to the first three days of priming. To generate a reference for passive dilution of a genomic nucleoside through DNA synthesis, we co-pulsed with an isotopically labelled 2’-deoxythymidine having a mass shift of 12 units ([^15^N_2_,^13^C_10_]-dT; hereafter referred to as dT+12; Fig. 2a). Due to its highly efficient incorporation, dT+12 was co-pulsed only during the last day of the m+4Met pulse. Thereafter, dT+12 was chased for two days in parallel to m+4Met. Remarkably, while in most cases the levels of modified cytosines readily declined with the onset of the chase, under a2i/LIF conditions hm+3dC and f+2dC accumulated for the first 12 and 24 hours, respectively (Fig. 2b). This shows again that the rate of Tet-mediated oxidation is quite high in mPSCs under a2i/LIF conditions and declines sharply upon priming, as already seen with the time course analysis during m+4Met labelling (Fig. 1b,c). Notably, under a2i/LIF conditions the decline of f+2dC after 36 h became quite steep, pointing to high turnover of fdC. Although under priming conditions oxidation of mdC to hmdC and further to fdC was hardly detectable in labelling time course experiments (Fig. 1), the fact that in the pulse chase experiments the decline of labelled hmdC and fdC was slower than that of dT+12 clearly shows that some mdC and hmdC oxidation still takes place (Fig 2b, central panel). Importantly, upon priming, the turnover of m+4dC is faster than the dilution rate of dT+12 (Fig. 2b, inset), revealing the presence of a component of the m+4dC turnover that cannot be explained by passive dilution through DNA synthesis.

**Figure 2.**
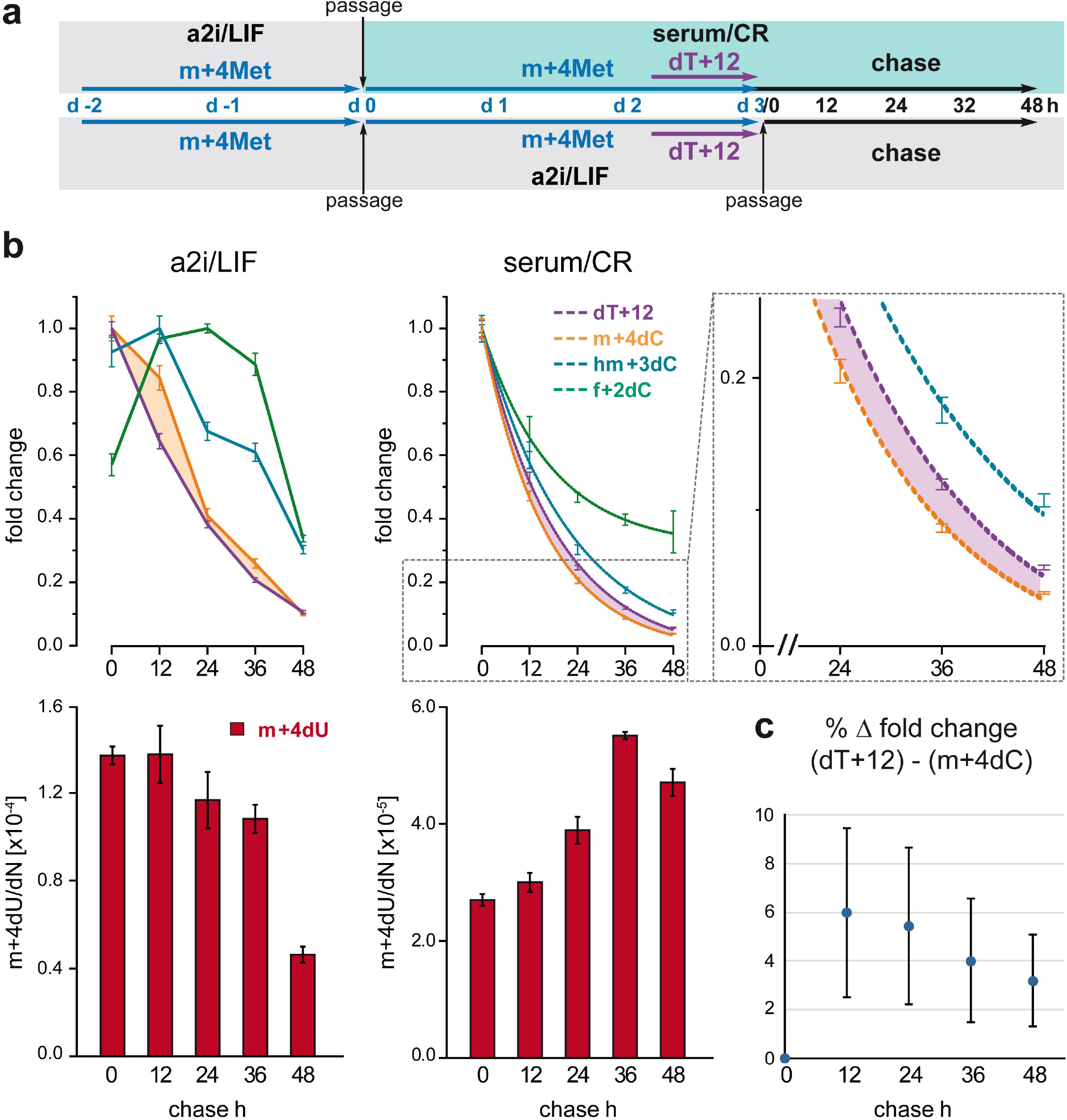
Turnover of genomic mdC and its derivatives under naïve and priming conditions. **a**) Workflow of m+4Met and dT+12 pulse-chase experiments with wild type mPSCs under naïve and priming conditions. **b**) Global profiles of the indicated labelled cytosine derivatives and dT+12 in the genome of naïve (a2i/LIF; left) and primed mPSCs (serum/CR; right and inset) upon chase of m+4Met. Upper panels show values relative to the highest level for each modification. Dashed lines represent first order decay fitted curves. Lower panels show absolute values of genomic m+4Met (note the different scales for m+4dU in the two conditions). Mean values and standard deviation of triplicate measurements are shown. **c**) Percent of difference between fold changes of dT+12 and m+4dC in wt mPSCs under priming conditions. Shown are mean and standard deviation of three independent biological replicates.

To assess the contribution of Tet-dependent oxidation to this component we performed the same pulse/chase experiment under priming condition (Fig. 2a, upper part) with E14tg2a mPSCs lacking all three Tet proteins (Tet triple knockout, alias Tet TKO)^35^ as well as parental, Tet 1-3 proficient E14tg2a cells (Fig. 3; note that E14tg2a cells are Hprt null^36^). In the parental cell line (Tet1-3 proficient), turnover of genomic m+4dC was again clearly faster than passive dilution of dT+12. After averaging the data from three such pulse chase experiments with wt mPSC lines, a paired two-tailed T-test showed high significance (P = 0.0053) for the deviation between m+4dC and dT+12 decay rates. This accounts for an active component of global mdC turnover involving 3-6% of the methylome, depending on the time point (Fig. 2c). In contrast, in the genome of Tet TKO cells the decline rate of m+4dC was essentially indistinguishable from that of dT+12 (Fig. 3). This shows that the active component of the mdC turnover is indeed largely dependent on the presence of Tet enzymes, and thus mostly due to mdC oxidation.

**Figure 3.**
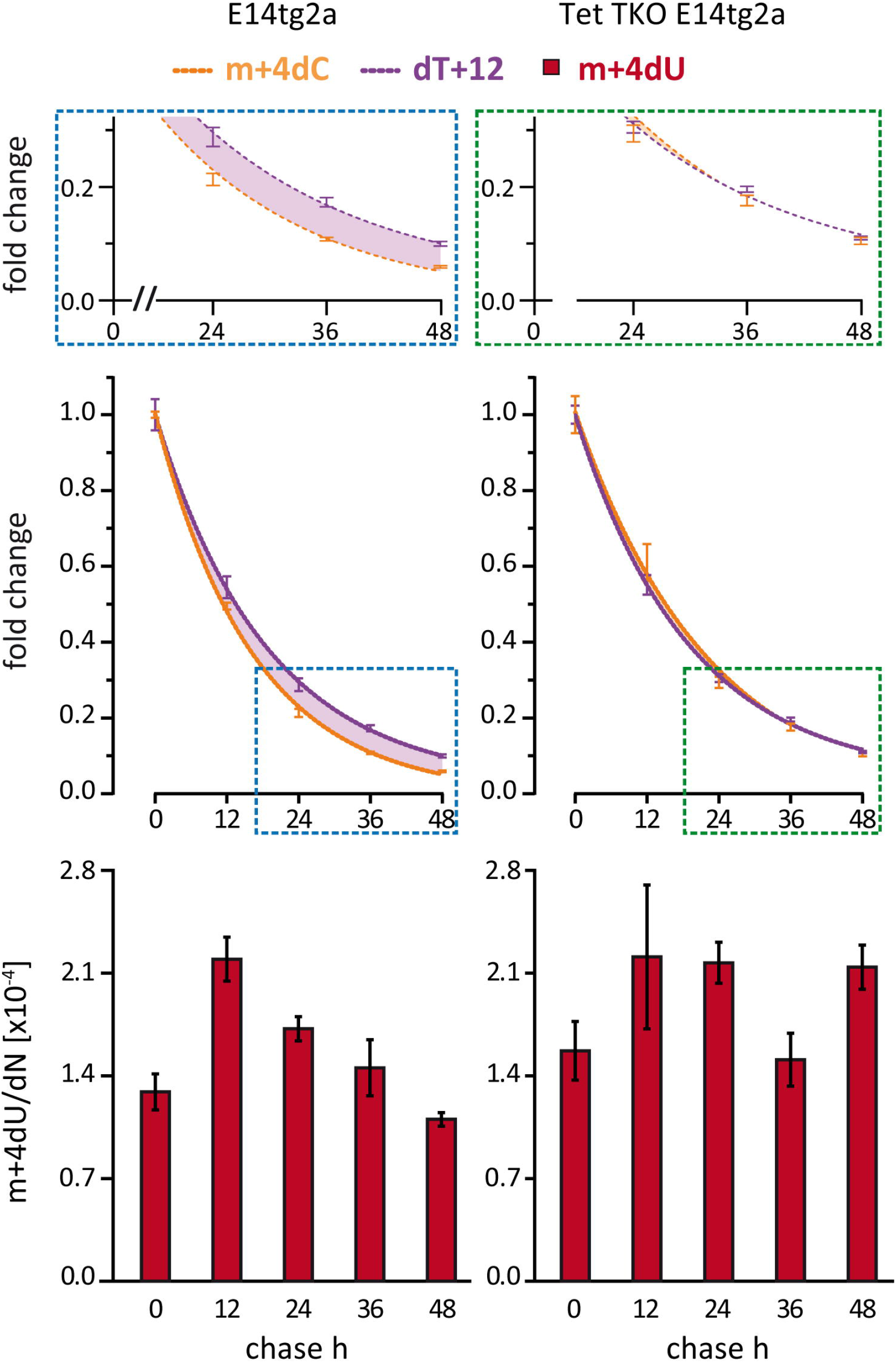
The active component of mdC turnover in primed pluripotent cells largely depends on Tet proteins. m+4Met and dT+12 pulse-chase experiment with parental and Tet TKO E14tg2a mPSCs under priming conditions (serum/CR). Profiles of global genomic levels of the indicated labelled cytosine derivatives and dT+12 are shown. Upper and mid panels show m+4dC and dT+12 values relative to the highest data point for each modification. Upper panels display magnifications of mid panels fields in dotted frames. Dashed lines represent first order decay fitted curves. Lower panels display global genomic levels of m+4dU.

### Additional mdC turnover through a DNA repair process

Next to oxidative mdC turnover, we started to investigate a potential deamination based pathway. Deamination of mdC and hmdC to dT (mdU) and hmdU, respectively, has been proposed as a potential alternative first step for active erasure of genomic C methylation. Upon metabolic labelling with m+4Met, deamination of m+4dC would generate m+4dU, that is dT bearing the m+4 methyl group (Fig. 1a). In order to investigate this process quantitatively we synthesised the isotopologue dT-[^13^C_5_,^15^N_2_] (Fig. 1b) and used it as a standard for UHPLC-MS^2^ analysis. As the source of methyl group for *de novo* dT biosynthesis is serine through 5,10-methylenetetrahydrofolate and not methionine through SAM, it is expected that, upon supplementation of the medium with m+4Met, m+4dC is the only source of m+4dU. To verify this expectation, we made use of mPSCs lacking all catalytically active Dnmts (Dnmt TKO), and therefore devoid of genomic mdC^37^. After priming in the presence of m+4Met, m+4dU was clearly detected in the genome of parental wt J1 mPSCs, but not in Dnmt TKO cultures. Thus, generation of m+4dU strictly requires the presence of m+4dC, showing that m+4dU originates exclusively through deamination of m+4dC (Supplementary Fig. S4).

Direct deamination of genomic mdC would imply the formation of T:G mismatches, which, if not efficiently repaired, would be highly mutagenic. It would then be reasonable to expect that a mechanism for highly efficient coupling of mdC deamination with BER of the ensuing T:G mismatches would have co-evolved^19^ so as to minimize the steady state levels of such mismatches. In contrast to this idea, we surprisingly detected substantial accumulation of m+4dU in genomic DNA, whenever cultures were provided with m+4Met (gDNA; Fig. 1–5, Supplementary Fig. S3-4, S7). To explain this observation we considered an alternative scenario whereby genomic strands containing m+4dC may first be subject to a DNA repair process leading to the release of soluble m+4dC/m+4dCMP, either by direct exonuclease action, as upon MMR and end resection of double strand breaks, or through digestion of excised strands (e.g., in nucleotide excision repair). Deamination of this material by Cytidine and Deoxycytidylate deaminases (Cda and Dctd)^38^, would generate soluble m+4dU/m+4dUMP, which could be phosphorylated and re-incorporated into the genome through DNA synthesis (Fig. 4a). To test if the high levels of m+4dU detected in the genome are generated along this pathway, we supplemented the medium with m+4Met and increasing concentrations of unlabelled dT. If DNA repair generates soluble m+4dU/m+4dUMP, incorporation of the latter into gDNA will be outcompeted by unlabelled dT. In addition, a dT excess will lead to high concentrations of dTMP and dTTP, which are both known to feedback negatively on Dctd through product inhibition and allosteric regulation, respectively^39^. The accumulation of genomic m+4dU in priming mPSC cultures was indeed progressively reduced with supplementation of increasing concentrations of unlabelled dT (Fig. 4b). This result supports the idea that genomic m+4dU originates in large part from a pool of soluble precursor.

**Figure 4.**
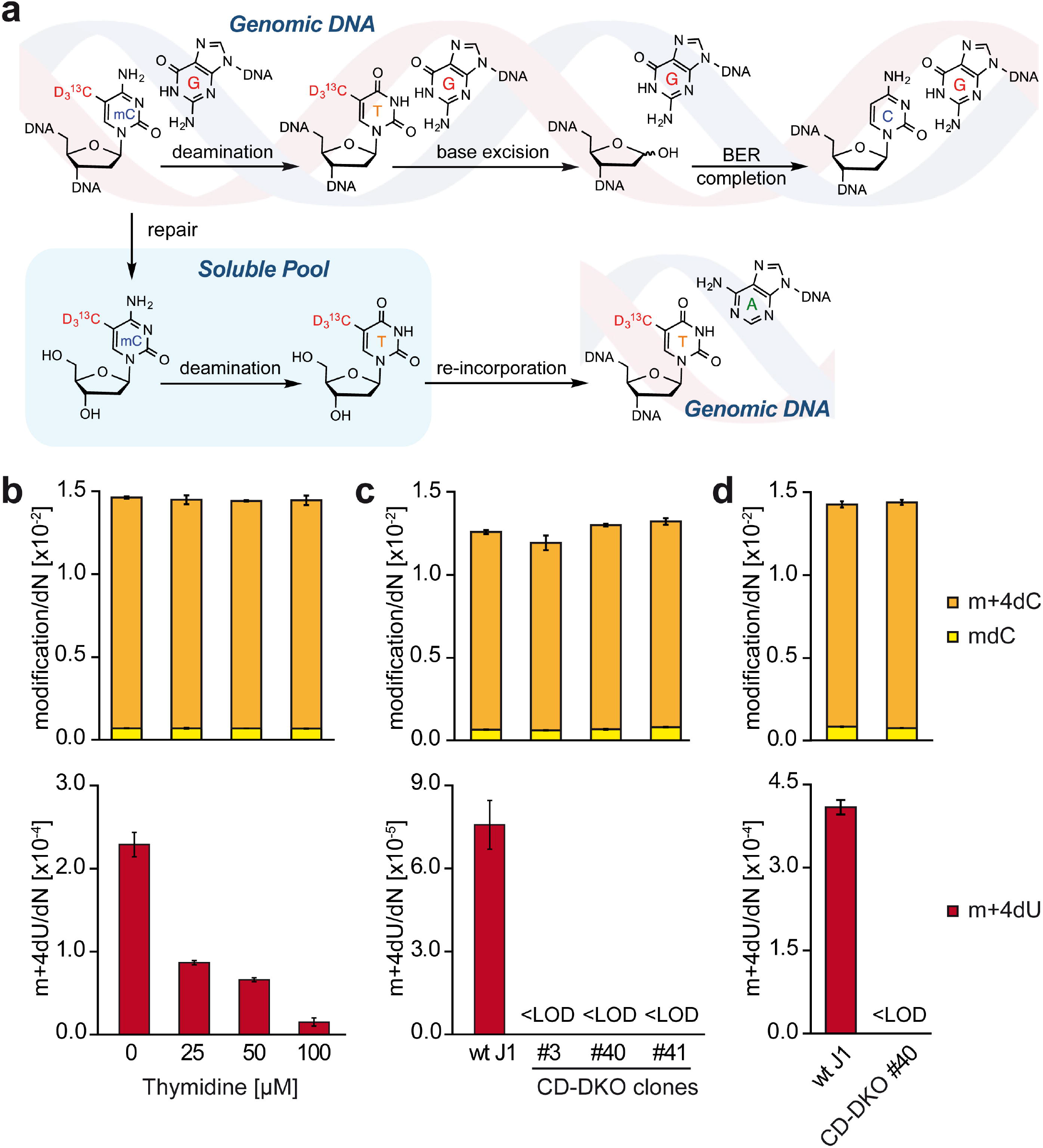
All detectable mdC deamination takes place in the soluble pool after DNA repair. **a**) Potential pathways for the conversion of genomic m+4dC into genomic m+4dU. **b**) wt mPSCs were primed under CHIR/IWR1 (serum/CR) conditions in the presence of m+4Met and growing concentrations of unlabelled dT in the culture medium as indicated. **c**) wt and CD-DKO clones mPSCs were grown under serum/LIF conditions in the presence of m+4Met. **d**) wt and CD-DKO clone #40 mPSCs were metabolically labelled with m+4Met under serum/CR priming conditions. In b-d priming and labelling was for five days. Global genomic contents of mdC, m+4dC (upper panels) and m+4dU (lower panels) are shown as mean and standard deviation of three technical replicates. LOD = limit of detection.

**Figure 5.**
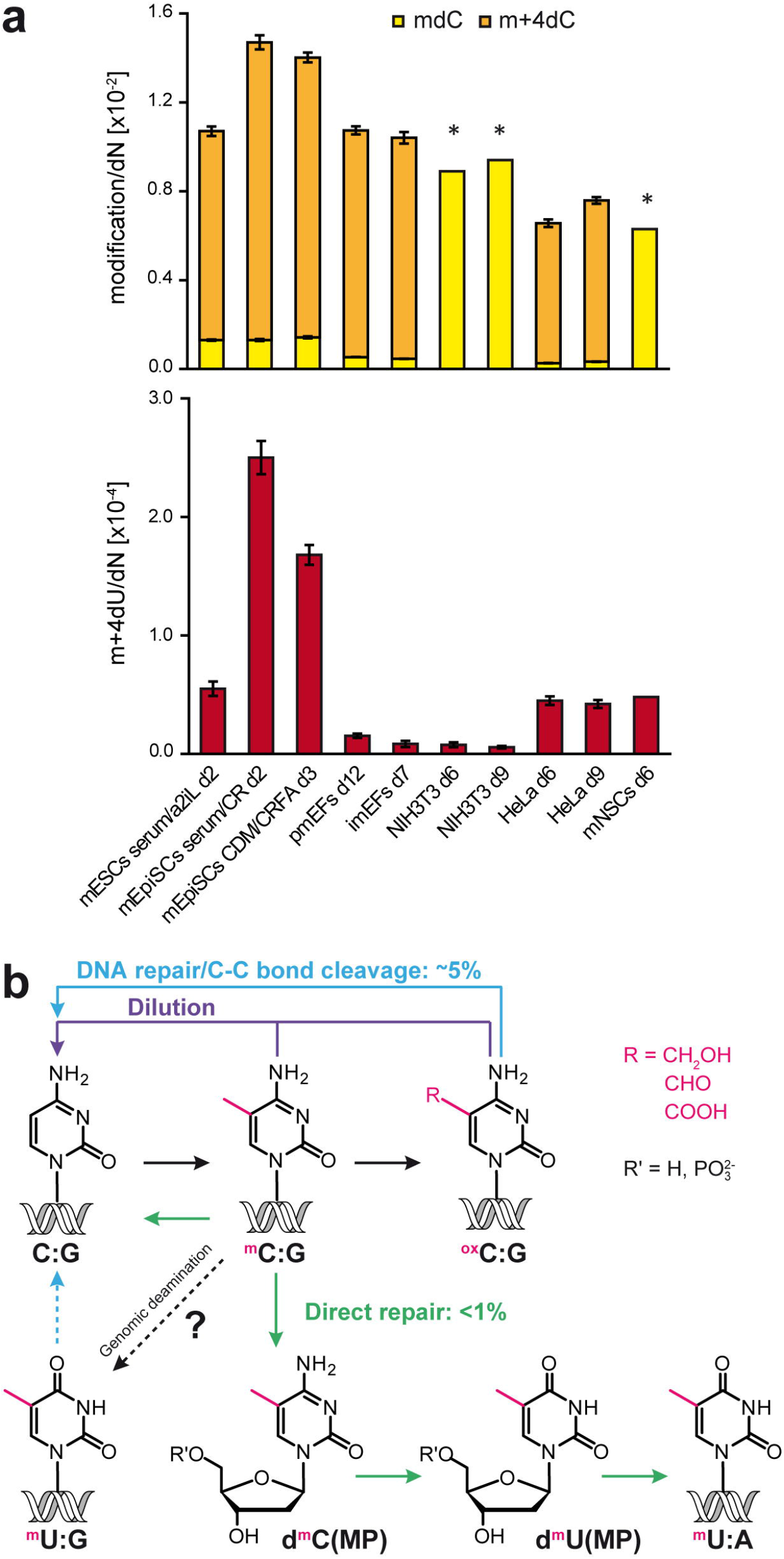
mdC turnover through oxidation-independent DNA repair is developmentally regulated. **a**) Cultures of mPSCs in naïve (a2iL = a2i/LIF) and primed states (CR = CHIR/IWR1; FA = FGF-2/Activin A; CDM = Chemically Defined Medium), primary (pmEFs) and immortalized mouse embryonic fibroblasts (imEFs and NIH3T3), HeLa and mPSC-derived neural stem cells (mNSCs) were labelled for the indicated number of days (d) with m+4Met. Asterisks indicate cases where distinct parallel cultures in the absence and presence of m+4Met were used to measure global levels of genomic mdC and m+4dU, respectively. In all other cases mdC, m+4dC and m+4dU were measure in the same culture labelled with m+4Met. Mean and standard deviation of three technical replicates are shown. **b**) Schematic summary of the dC modification turnover pathways identified in primed pluripotent cells. Deamination of genomic mdC followed by BER could not be detected and may occur only at levels below the sensitivity of our assay.

To further probe whether deamination of m+4dC takes place within gDNA or at the level of soluble nucleoside/nucleotide pools, we next used CRISPR/Cas9-mediated gene editing to generate mPSC with double deficiency for Cda and Dctd (CD-DKO; Supplementary Fig. 5a,b). Genomic sequencing showed that three of the isolated clonal lines bear compound biallelic *Cda* and *Dctd* mutations incompatible with the expression of functional enzymes (Supplementary Fig S5c-e). All three CD-DKO clones grew at slower rates than parental wt cells, a phenotype which was at least partially relieved by supplementation of the medium with dT (Supplementary Fig. S5f). This indicates that deamination of (d)C/(d)CMP by Cda and/or Dctd provides substantial amounts of (d)U/(d)UMP for biosynthesis of (d)T/(d)TMP by Thymidine synthase to support DNA replication and is fully consistent with previously reported dT auxotrophy of Dctd deficient somatic cell lines^40^. As two of the CD-DKO clones grew particularly poorly under serum/CR priming conditions, we first used serum/LIF medium for metabolic labelling with m+4Met under priming conditions. Importantly, while the genomic content of mdC/m+4dC was the same as in parental wt J1 cells, only background levels of m+4dU could be measured in all three CD-DKO clones (Fig 4c). The same result was obtained with CD-DKO clone 40 upon priming in serum/CHIR/IWR1 medium (Fig. 4d). These results clearly show that the genomic m+4dU measurable in wt cells with our UHPLC-MS^2^ method originates from a process involving direct repair of gDNA containing m+4dC, with consequent release of soluble m+4dC/m+4dCMP, followed by deamination to m+4dU/m+4dUMP and (re)-incorporation of the latter into the genome through phosphorylation to the triphosphate state and DNA synthesis. Although our data point to exclusive deamination in the soluble pool, we wanted to further investigate potential deamination events of mdC to T directly in the genome, particularly in light of previous reports in this direction^17,19^. In this respect it is important to consider the limitations of our quantitative UHPLC-MS^2^ method. In UHPLC the overwhelming amount of natural dT present in the genome co-elutes with m+4dU and potentially outcompetes it during ionization for the capture of charge. The effect is a strong ion suppression, resulting in a reduction of the m+4dU signal and, consequently, a high limit of detection (LOD). To put this effect into perspective we calculated that our LOD for m+4dU is about 100 times higher than that for fdC (around 1×10^−5^ versus 1×10^−7^ per nucleotide), which is a relatively stable dC modification considered to have an epigenetically relevant role.

However, deamination of very few m+4dC residues directly within the genome could trigger long patch BER and/or MMR^20,21^, leading to the release of much larger amounts of m+4dC in the soluble pool and, subsequently, deamination to m+4dU and re-incorporation into the genome, explaining the large amounts of genomic m+4dU observed. In CD-DKO cells direct deamination of genomic m+4dC would be expected to be the only potential source of m+4dU and, as the latter would then be mismatched to G, its steady state levels would be expected to be very low due to efficient repair. In order to probe genomic deamination with higher sensitivity the ion suppression effect had to be substantially reduced. To this end we performed a double labelling experiment with m+4Met and a dC nucleoside where 9 hydrogen atoms are replaced with deuterium (dC[D_9_]; Supplementary Fig. S5a). After incorporation into the genome, dC[D_9_] will be methylated to m+4dC[D_8_], which deaminates to m+4dU[D_8_] (a dT with eleven D atoms and a +12 mass shift; Supplementary Fig S6a). This massive H-to-D exchange was chosen as it is known to determine a shift to shorter retention time in UHPLC, allowing m+4dU[D_8_] to escape co-elution with dT. Indeed, co-injection tests showed a slightly shorter retention time for dT[D_8_] relative to dT (Supplementary Fig. S6b). Notably, m+4dU[D_8_] is expected to have an even shorter retention time that would allow it to at least partly escape ion suppression from natural dT. After double labelling with m+4Met and dC[D_9_] we were able to detect m+4dU[D_8_] confidently only in wt mPSCs (Supplementary Fig. S6c). However, we sporadically detected small m+4dU[D_8_] peaks in technical replicates of CD-DKO samples. After integration of these sporadic signals, the corresponding values were invariably below LOD, defined as three times the average level of blank samples. Although we cannot exclude that these signals are caused by background processes, we nevertheless performed a calculation taking into account the limits of detection and quantification that were gained by deuterium labelling. This calculation allows us to estimate the maximum number of potential genomic deamination events between 500 and 5000 m+4dU[D_8_] molecules per cell in the course of our labelling experiment. According to an estimated rate constant for spontaneous hydrolytic deamination of mdC in solution^41^ only around 20 such events would be expected to take place. Therefore, although a reliable and reproducible signal for m+4dU[D_8_] could not be detected in CD-DKO cells, we cannot exclude the occurrence of genomic deamination above the rates of spontaneous hydrolysis. However, in the best case this process cannot exceed a level of 1×10^-6^-1×10^-7^ events per nucleotide.

### Tet enzymes do not trigger the DNA repair process responsible for turnover of genomic mdC into dT

Conversion of m+4dC into m+4dU obviously does not require oxidized intermediates. However, it was recently suggested that in the zygote Tet3 generates undefined DNA lesions, triggering repair-mediated erasure of genomic methylation from the paternal genome^42^. Also, hmdC may mark DNA damage sites and promote their repair^43^ or even target DNA repair by selectively recruiting endonucleases like Srap1^44^ and Endonuclease G^45^. We therefore considered whether Tet proteins contribute to the turnover of genomic mdC into dT by generating hmdC or low levels of DNA lesions such as hmdU^27^, which triggers BER initiated by the DNA glycosylase Smug1. To this aim we primed Tet TKO and parental E14tg2a mESCs in the presence of m+4Met. Surprisingly, higher levels of genomic m+4dU were detected in the absence of Tet enzymes (Fig. S7). Similarly, analysis of the pulse chase experiment with the same pair of cell lines also revealed a tendency for higher accumulation of m+4dU in Tet TKO mPSCs at later chase time points (Fig. 3, lower panels). Therefore, in pluripotent cells Tet proteins do not seem to trigger DNA repair events that contribute to turnover of mdC into dT.

### Oxidation-independent repair of mdC-containing gDNA is developmentally regulated

To investigate whether oxidation-independent mdC turnover is regulated during the transition to primed pluripotency, we first compared the genomic abundance of m+4dU in the time course and pulse-chase experiments under naïve and priming conditions described above (Fig. 1–3, Supplementary Fig. S3). In time course experiments, progressive accumulation m+4dU was much faster under priming conditions, reaching approximately five times higher levels than under naïve conditions (Fig. 1, Supplementary Fig. S3). Also, m+4dU was the only m+4dC derivative that accumulated during label chase under priming conditions, though to different extents in the two cell lines tested (Fig. 2b,3). Thus, in contrast with the observed decline of mdC oxidation (Fig. 1,2, Supplementary Fig. S3), these results reveal increased rates of oxidation-independent repair of mdC-containing gDNA upon acquisition of primed pluripotency. This anti-correlation suggests that oxidation-independent mdC turnover through DNA repair may serve at least in part as a supplementary pathway for the erasure of dC methylation. However, as in primed Tet TKO mPSCs there was no apparent difference between global mdC turnover and passive dilution through DNA synthesis (Fig. 3: decay of m+4dC and dT+12, respectively), this process cannot involve more than 1% of the methylome.

At last, we compared oxidation-independent mdC turnover through DNA repair in pluripotent states (intermediate serum/a2i/LF as well as primed serum/CR and EpiSCs under CDM/CHIR/IWR1 /FGF-2/Activin A conditions) with various lineage committed cell types, including primary and immortalized embryonic fibroblasts, neural stem cells (NSCs)^46^ and HeLa cells. Although primed pluripotent cells were labelled with m+4Met for only two to three days, m+4dU reached higher levels in their genome than in those of any of the somatic cell types cultured for substantially longer periods, ranging from 6 to 12 days (Fig. 5a). This clearly shows that in lineage committed cells oxidation-independent mdC turnover through DNA repair is substantially lower than in primed pluripotent cells. In addition, the rate of oxidation-independent mdC turnover is clearly not proportional to the global genomic abundance of mdC either between pluripotent states or among pluripotent and lineage committed cells. In particular, among the latter oxidation-independent mdC turnover is clearly more active in NSCs than in fibroblasts, despite the mdC genomic content being at least 20% lower in NSCs. These observations indicate that oxidation-independent mdC turnover is under developmental and cell lineage-dependent control.

## Discussion

Recent methylome analysis of matched clonal and polyclonal cell populations^47^ and studies on the contribution of Dnmt3 proteins to DNA methylation maintenance^48,49^ suggested that in heterogeneously primed mPSCs cultures (serum/LIF) genomic mdC undergoes higher dynamic turnover as compared to differentiated somatic cells. However, the extent and especially the nature of this turnover have not been addressed. Previously, Bachmann *et al*. reported pulse-chase experiments with m+4Met to estimate the global turnover of genomic dC modifications in mPSCs cultured under naïve conditions (2i/LIF), using a very short pulse time^50,51^. However, the naïve pluripotent state is characterised by very low global levels of genomic dC modifications, which increase only upon exit from the naïve state and acquisition of primed pluripotency^1,23^, a crucial transition to lay the ground for commitment to somatic lineages.

Here, we report the first analysis of on- and off-rate kinetics for genomic dC modifications upon transition to primed pluripotency. We show that in mPSCs under a2i/LIF conditions, which display expression and genomic methylation patterns that are intermediate between naïve and primed pluripotent states and resemble those reported for the formative state^33^, the rates of mdC oxidation to hmdC and further to fdC are quite sustained and the removal/conversion of fdC is very rapid. This points to a very dynamic equilibrium for fdC under these conditions. It is tempting to speculate that these high levels of oxidative turnover may underlie the frequent periodical oscillations in global mdC levels recently identified through single cell analysis in partially (and heterogeneously) primed serum/LIF cultures^52^. In contrast to the high levels of mdC oxidation in a2i/LIF conditions, upon transition to a more advanced, primed pluripotency state the global oxidation rates of mdC and hmdC as well as the removal of hmdC and fdC decline drastically. Despite this the global turnover rate of genomic mdC is 3-6% faster than can be accounted for by passive dilution through DNA synthesis (DNA replication and repair; Fig. 5b) and this difference is largely due to Tet-mediated mC oxidation. Evidence for passive erasure of dC methylation based on impairment of its maintenance has been provided for primordial germ cells, preimplantation embryonic development and the reversion from heterogeneous partially primed/formative states present in serum/LIF cultures to the naïve pluripotent state^1,24^. We suggest that against the backdrop of global gain and maintenance of dC methylation occurring in the forward developmental transition from naïve to primed pluripotency an active demethylation mechanism may be more attainable than preventing maintenance of dC methylation at selected sites, especially in the case of loci with a highly accessible chromatin conformation like actively transcribed genes.

Furthermore, by tracing the origin of genomic m+4dU to deamination of soluble m+4dC, we show that a small part of genomic mdC is turned over by a DNA repair process that involves its direct excision from the genome (Fig. 5b). While we show that TET proteins are clearly not involved in triggering or targeting this non-oxidative mdC turnover, our data neither support nor rule out that low levels of direct enzymatic deamination of genomic mdC may trigger the DNA repair process underlying this turnover pathway. Although the nature of this DNA repair process and the events triggering it remain to be determined, its rate is not proportional to global genomic mdC levels, either between pluripotent states or between lineage committed cells, indicating that the underlying DNA repair process is developmentally regulated. In this regard, it should be noted that in PSCs, including primed human PSCs, the expression of DNA repair factors and the proficiency of various DNA repair pathways were shown to be higher than in somatic cells^53–55^. Interestingly, the extent of oxidation-independent turnover of mdC through DNA repair is not proportional to global genomic mdC levels, either between pluripotent states or lineage committed cells, indicating that the underlying DNA repair process does not take place randomly throughout the methylome. In addition, the levels of oxidation-independent mdC turnover show an inverse correlation with mdC oxidation rates across pluripotent states and seem to be slightly more sustained in Tet deficient cells, pointing to a possible function as alternative or supplementary mechanism dedicated to active erasure of cytosine methylation. Whether this is indeed the case requires further investigation.

## Acknowledgements

We are very thankful to the following colleagues: Markus Möser (Max Planck Institute for Biochemistry, Martinsried, Germany) for K3 mPSCs (Kindlin3+/+); Hitoshi Niwa and Masaki Okano (both at Kumamoto University, Japan) for Oct4-YFP-Puro mPSCs and Dnmt TKO J1, respectively; Yi Zhang (Boston Children’s Hospital, Boston, MA) for parental and Tet TKO E14tg2a. Anti-DCTD antibody and purified recombinant DCTD protein were generous gifts from Frank Maley (New York State University, NY), Sebastian Bultmann and Christopher Mulholland for guidance on high depth bisulfite amplicon sequencing (both at Ludwig Maximilian University, Munich).

Funding was provided by the Deutsche Forschungsgemeinschaft via the programs SFB1309 (TP: A4), SFB1361 (TP: 2), GRK 2338 and SPP-1784. This project has received funding from the European Research Council (ERC) under the European Union’s Horizon 2020 research and innovation programme (grant agreement n° EPiR 741912). YZ is supported by the China Scholarship Council (CSC Nr. 201806200069).

## Author Contributions

FS and TC conceived the study and designed experiments. FS, SS, AK, YZ, GA, OK and JS performed experiments and analyzed data. FS, TC and SS interpreted data. RR, CE and EK synthesized isotopically labelled nucleosides used as standards for LC-MS/MS. FS, TC and SS wrote the manuscript.

## Data availability and code availability statements

The raw data that support the findings of this study are available from the corresponding authors upon reasonable request. No own code was used.

## Methods

### Cell culture

Medium components were from Sigma unless specified otherwise. Basal medium for mPSC culture was DMEM high glucose containing 10% FBS (Pan-ES, Pan-Biotech), 2 mM L-Alanyl-L-Glutamine, 1x MEM Non-essential Amino Acid Solution and 0.1 mM ß-mercaptoethanol. mPSC lines were routinely maintained in basal medium supplemented with 1000 U/mL LIF (ORF Genetics), 3 μM CHIR99021 and 7.5 μM CGP77675 (a2i). For adaptation to 2i/LIF conditions CGP was replaced with 1 μM PD0325901. For priming, mPSC basal medium was supplemented with 1.5 μM CHIR99021 and IWR1-endo at 2.5 μM as previously reported^26,56^. For experiments under serum/LIF conditions basal medium was supplemented exclusively with 1000 U/ml LIF. Small molecule inhibitors were purchased from Selleckchem (CHIR, PD, IWR1), Axon Medchem (CHIR, CGP, PD, IWR1) or MedChemExpress (CHIR). Gelatin coating was used for all mPSCs cultures in serum containing medum, except for PR8 cultures where coating with 2.2 μg/cm^2^ Laminin 521 (BioLamina) was necessary. For metabolic labelling experiments with m+4Met under all conditions mPSC basal medium was generated with 0.2 mM L-Methionine- and L-Cysteine-free DMEM (Sigma or Life Technologies) supplemented with m+4Met ([*methyl*-^13^C,d_3_]) and L-Cysteine. EpiSCs in chemically defined serum-free medium (CDM) were derived from the Oct4-YFP reporter mPSC line OLY2-1^25^. These were first adapted to CDM supplemented with CHIR, PD and LIF as reported above, then transferred to CDM supplemented with CHIR, CGP, 12 ng/ml FGF-2 (Miltenyi Biotec) and 20 ng/ml Activin A (PeproTec) and maintained in the latter medium by passaging as small cell clusters using 5 mM EDTA in Hank’s balanced salt solution without Mg^2+^, Ca^2+^ and sodium carbonate supplemented with 10 mM HEPES buffer. CDM was DMEM high glucose supplemented with 1x NEAA, 2 mM L-Alanyl-L-Glutamine, 15 μg/ml human recombinant Insulin, 10 μg/ml human holo-Transferrin (Merck), 12.5 mg/ml AlbuMAX I (Life Technologies) 0.7 μg/ml vitamin B12, 1.8 ng/ml biotin, 0.75 μM ZnSO_4_ and 2.6 nM CuSO_4_. Laminin 521 coating was used for all mPSC cultures in CDM. The neural stem cell line ENC1 was generated and cultured as previously described^46^. ENC1 cells were labelled with m+4Met ([*methyl*-^13^C,d_3_]) in DMEM/F-12 generated with L-Methionine-free DMEM supplemented with N2^57^ and 20 ng/ml each of FGF-2 and EGF (PerpoTech). Primary mEFs (CF-1, Applied StemCell) were cultured in mPSC basal medium. Somatic cell lines were cultured in DMEM containing 10% FBS (Life Technologies), 2 mM L-glutamine and 0.1 mM β-mercaptoethanol.

### Generation of CD-DKO mPSCs by CRISPR/Cas9 editing

Single guide RNAs were designed over exon1 and 4 of Cda and Dctd, respectively (Fig. S5), using CRISPR Design Tool (http://crispr.mit.edu/). Oligonucleotides used for construction of Cas9/gRNA expression vectors are listed in Supplementary Table 1. The oligonucleotides were annealed and cloned in the BbsI site of pSpCas9-2A-Puro (PX 459; Addgene Plasmid 48139)^58^. Cas9/gRNA expression vectors targeting both Cda and Dctd were co-transfected in wt J1 mPSCs using Lipofectamine 2000 (Life Technology) according to the manufacturer’s instructions. Two days after transfection cultures were selected with 1 μg/ml puromycin for two days and subcloned by limiting dilution. Clones were screened using the Surveyor Mutation Detection Kit (Transgenomics) and potential compound biallelically targeted clones were subject to sequencing. For reverse transcription-PCR, RNA was isolated with the ZR-Duet DNA/RNA MiniPrep Kit (Zymo Research) and first strand cDNA synthesis was as previously described^59^. Primers were used to amplify the Cda cDNA are reported in Supplementary Table 1. For western blot analysis of Dctd whole cell extracts and purified human recombinant DCTD protein^60^ were probed with a rabbit anti-DCTD antibody^39^.

### Isolation of genomic DNA and UHPLC-MS^2^ analysis

Isolation, digestion and UHPLC-MS^2^ analysis of gDNA samples were performed as described in Traube et al.^61^ with the following modifications: for every technical replicate 4 μg of gDNA were digested and analysed; the enzyme mixture including S1 Nuclease, Antarctic Phosphatase and snake venom Phosphodiesterase was used for digestion; the digestion mixture was supplemented with 0.8 μM Tetrahydrouridine (Abcam) to inhibit potential cytidine deaminase activity in enzyme preparations.

### Chemical synthesis

For a description of the synthesis of labelled nucleosides see Supplementary Methods in Supplementary Information file.

