## Supplementary Information for "Oxidative and non-oxidative active turnover of genomic methylcytosine in distinct pluripotent states"

**Supplementary Figures S1-S7**

**Supplementary Table 1**

**Supplementary Methods**

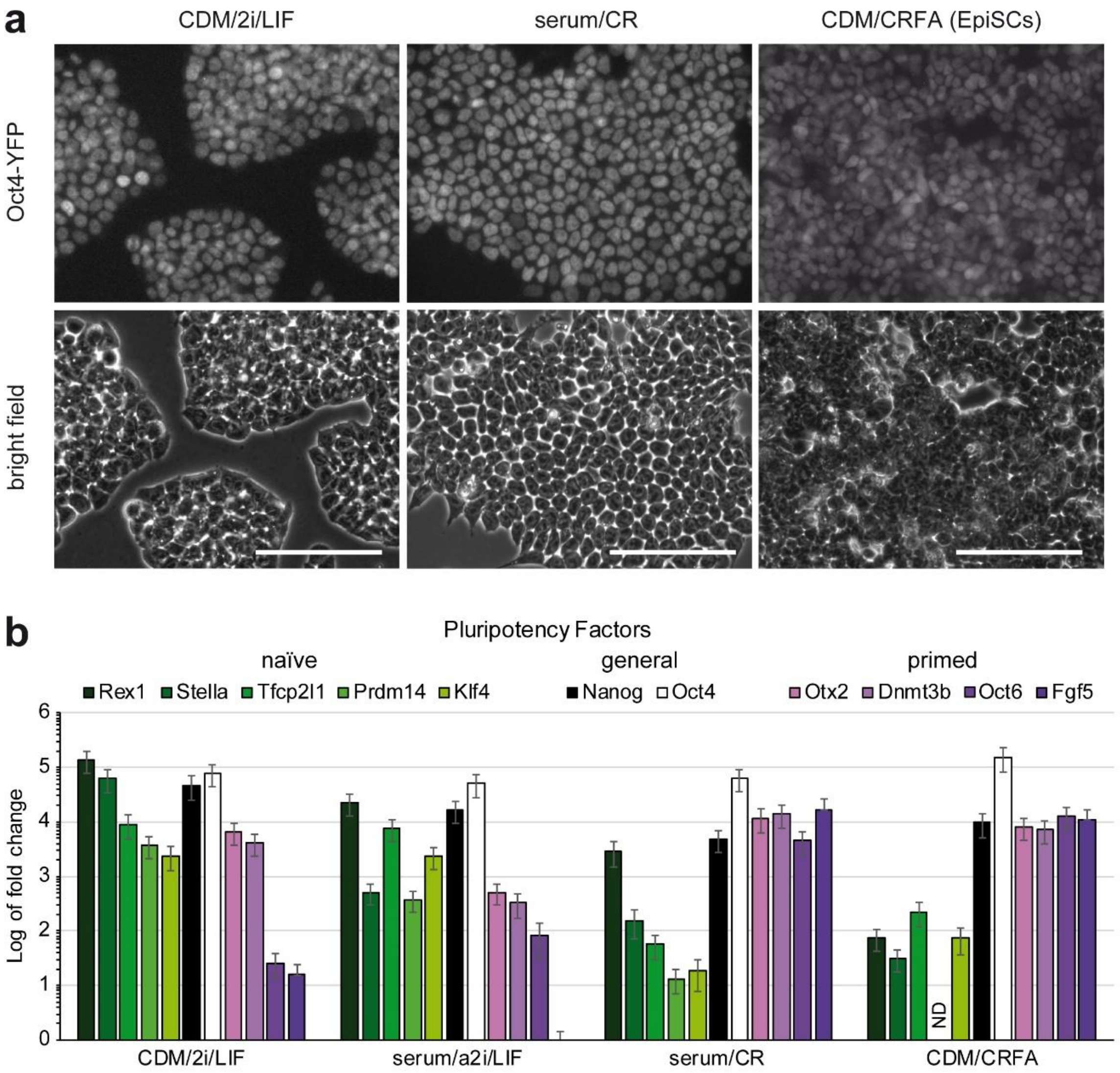

**Figure S1. Expression of pluripotency and early lineage specification factors in mPSCs under different culture conditions. a**) YFP fluorescence and bright field images of an Oct4-YFP reporter mPSC line after adaptation to naïve conditions in chemically defined medium (CDM/2i/LIF; left), followed by six days of priming under serum/CHIR/IWR1 conditions (serum/CR; middle) or permanent priming to mEpiSCs by passaging under chemically defined medium containing CHIR/IWR1/FGF-2/Activin A (CDM/CRFA; right). Scale bars = 100 µm. **b**) Transcript levels of pluripotency factors assayed by reverse transcription-qPCR on cultures of the Oct4-YFP mPSC line shown in a under CDM/2i/LIF (naïve), serum/a2i/LIF as well as both serum/CR and CDM/CRFA primed conditions. Note that under serum/a2i/LIF conditions relative high and low levels of naïve and primed pluripotency factors are expressed, respectively, while the levels of Oct4 and Nanog transcripts are similar to those in naïve conditions. Also, under transiently primed conditions (serum/CR) the levels of naïve and primed pluripotency factor transcripts are similar as in indefinitely primed EpiSCs under

CDM/CRFA conditions. Mean Log of fold change values and standard errors of three technical replicates are shown. ND = not detectable. Gapdh was used as house keeping gene reference to calculate ΔCt values and expression fold change values thereof.

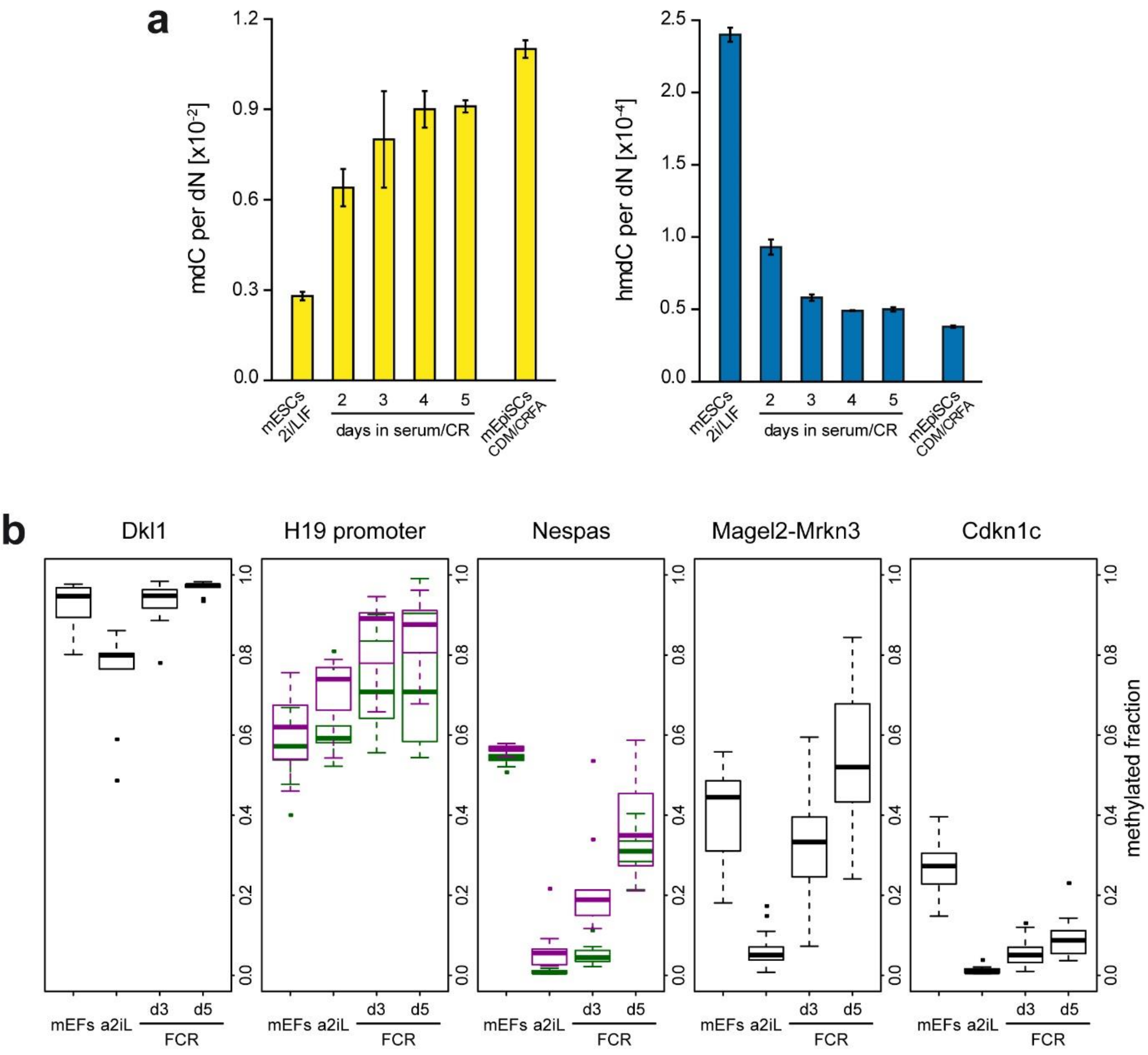

**Figure S2. Global and local transitions of genomic mdC and hmdC levels upon priming of mPSCs. a**) Global levels of genomic mdC (upper panel) and hmdC (lowr panel) assayed by UHPLC-MS/MS in wt J1 mPSCs transitioning from naïve state (serum/2i/LIF; day 0) to primed state over five days in serum/CHIR/IWR1 (serum/CR) and the Oct4-YFP reporter mEpiSC line shown in C derived and permanently cultured in serum-free CDM containing CHIR/IWR1/FGF-2/Activin A (CRFA). **b**) High depth targeted amplicon bisulfite sequencing analysis of selected secondary imprints in mPSCs upon transition from a2i/LIF to serum/CR conditions. Boxes display the inter-quartile range (IQR), where lower and upper edges represent the first and third quartile, respectively, and the thicker orizontal line is the median. Whiskers mark 3xIQR or 1.5xIQR where outliers (dots) below and above 3xIQR are present. Magenta and Green box plots represent distinct amplicons within the *H19* promoter region and *Nespas*. Primary mouse embryonic fibroblasts (mEFs) are shown as reference for differentiated primary somatic cells.

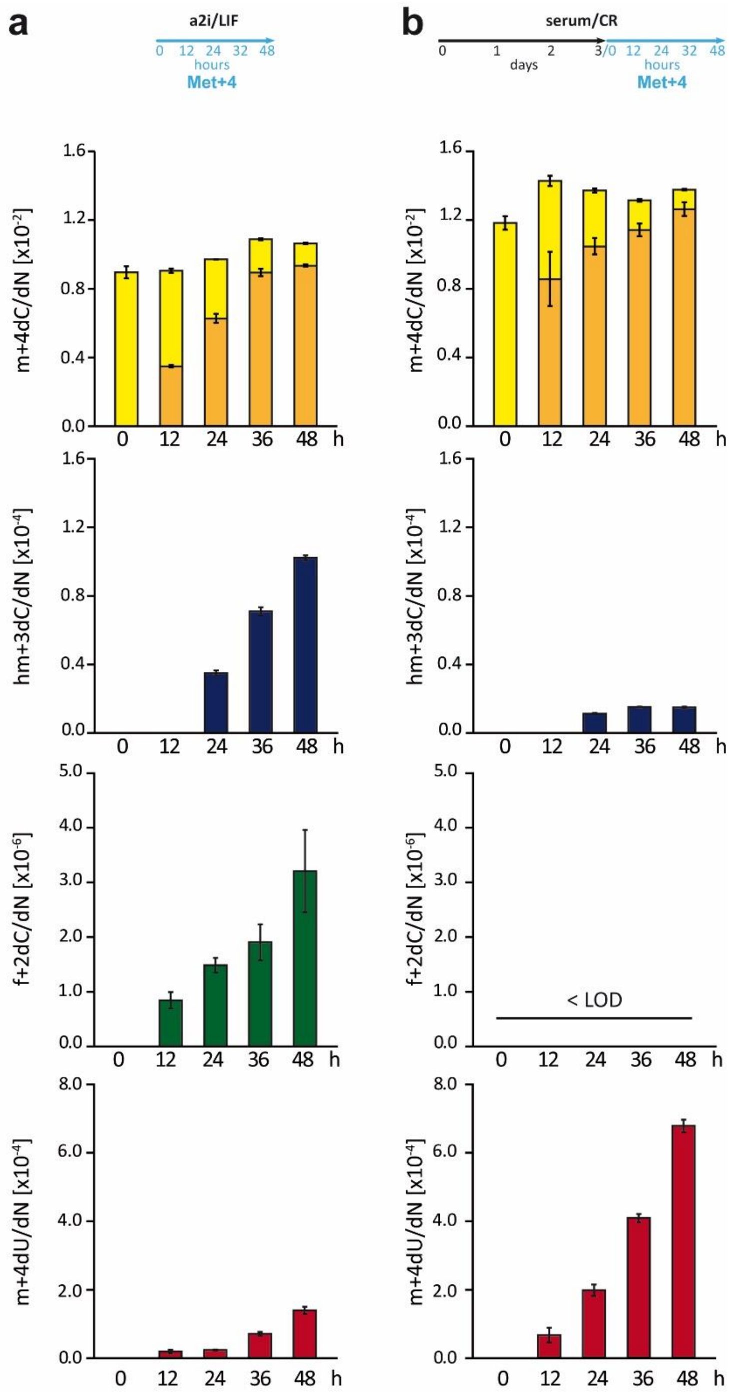

**Figure S3. Drastic reduction of global genomic mdC oxidation and increase of mdC to mdU conversion upon transition to primed pluripotency** (related to Fig. 1). Time course analysis of genomic dC derivatives upon metabolic labelling with m+4Met under naïve (**a**; a2i/LIF) and priming (**b**; serum/CR) conditions. The labelling/time course schedules are shown at the top. Global levels of unlabelled (pale yellow) $^{m}C$ and m+4dC (dark yellow), hm+3C (blue) and f+2C (green) are shown as mean and standard deviation of three technical replicates. Independent biological replicate of the experiment shown in Fig. 1b,c. LOD = limt of detection.

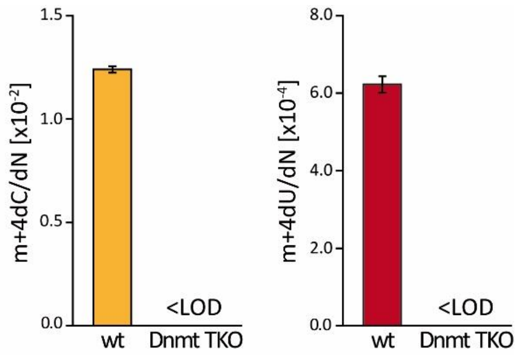

**Figure S4. Genomic m+4dU is generated exclusively from genomic m+4dC.**
Global levels of m+4dC (yellow) and m+4dU (red) in the genome of wt and Dnmt TKO J1 mPSCs after priming for five days in the presence m+4Met. In Dnmt TKO cells neither m+4dC nor m+4dU were above background levels (LOD = limit of detection). Mean and standard deviation of three technical replicates are shown.

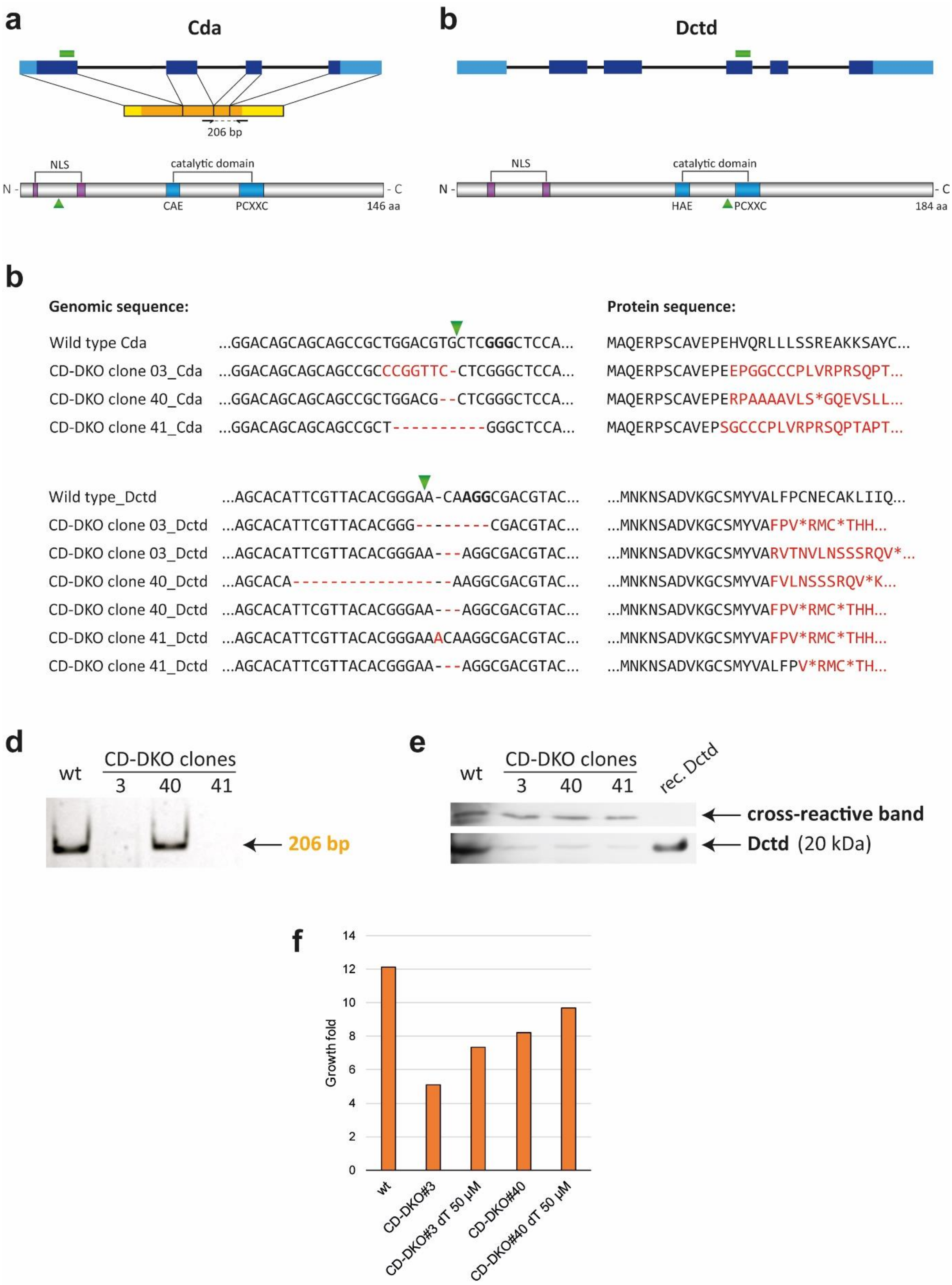

**Figure S5. Generation and characterization of CD-DKO mPSCs** (related to Fig. 4).

**a** and **b**) Schematic representations of gene body (top), transcript (middle, A only) and protein (bottom) for mouse Cda (a) and Dctd (b). Exons are represented by blue boxes, with UTR regions in lighter colour. The positions of gRNAs are indicated by green rectangles over the gene, while green triangles indicate the approximate positions of the expected truncation of the protein. The catalytic domains are shown along with the two $Zn^{2+}$ co-ordinating motifs, which are conserved across kingdoms of life among all cytosine nucleoside and monophosphate nucleotide deaminases, including the vertebrate-specific Aid/Apobec family. **c**) Genomic sequences of Cda and Dctd alleles as well as respective protein sequences for Cda and Dctd in CD-DKO clones. Antisense strands are shown so as to highlight the PAM sequences (bold type in wt sequence). The expected Cas9 cut sites are indicated by green triangles. The three clones are homozygous and heterozygous with respect to Cda and Dctd mutations, respectively. Mutations in Dctd were targeted to exon 4 and lead to termination of the frame before the second conserved catalytic motif. Note that, though exons 3 and 5 are in frame, alternative splicing events skipping the frameshifts in exon 4 would generate a protein lacking both conserved motifs involved in $Zn^{2+}$ co-ordination, which would therefore lack catalytic activity. **d**) Reverse transcription-PCR products from CDA transcripts in wt cells and CD-DKO clones. **e**) Western blot with an anti-Dctd antibody on whole cell extracts from wt and CD-DKO cells and purified recombinant Dctd protein. Cross-reactive bands are shown in the upper panel as loading control. As all the alleles detected in CD-DKO clones are expected to result in substantial protein truncations, the week signals visible in CD-DKO lanes at approximately the same position as in wt and recombinant Dctd lanes are likely due to unspecific cross-reaction. **f**) Growth delay of CD-DKO mPSCs and partial rescue with dT. The same number of cells was plated for two days under a2i/LIF conditions with or without addition of 50 µM dT to the medium (only CD-DKO clones). Cell counts after two days of culture are shown.

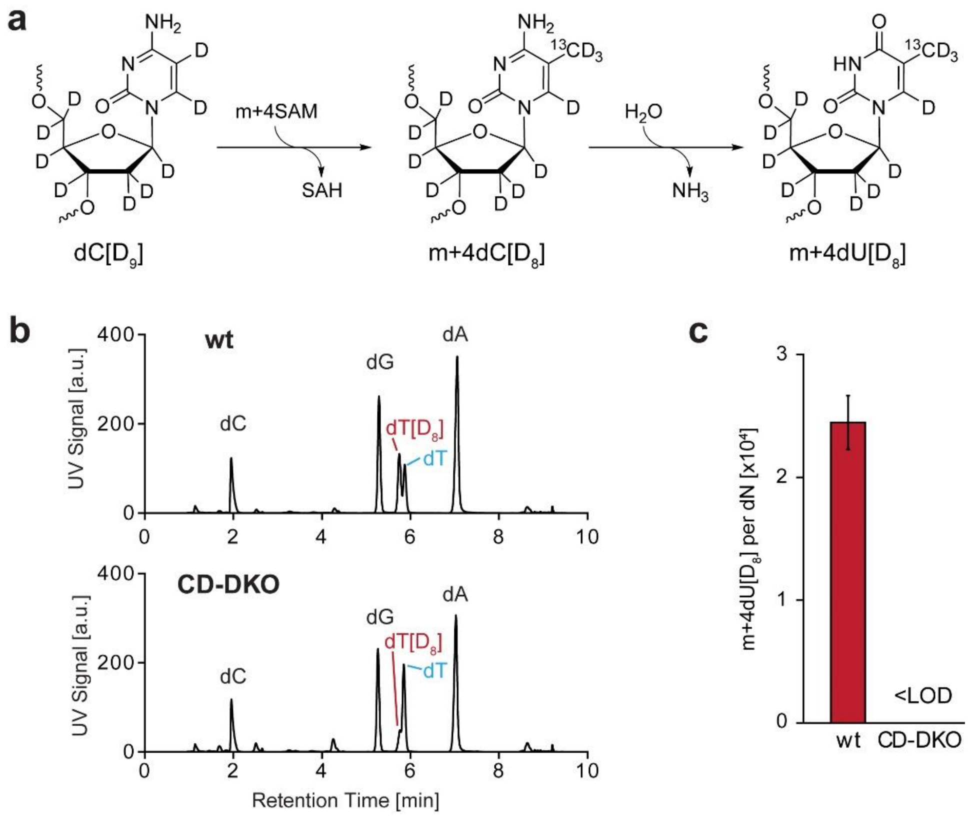

**Figure S6. Double labelling of primed wt and CD-DKO J1 with m+4Met and dC[D$_9$]** (related to Fig. 4).

**a**) Chemical structures and interconversion of dC[D$_9$], m+4dC[D$_8$] and m+4dU[D$_8$]. SAH = S-adenosylhomocysteine. **b**) UHPLC chromatogram (UV trace) showing the retention delay of dT[D$_8$] relative to natural dT. **c**) Quantification of m+4dU[D$_8$] accumulated in wt and CD-DKO PSCs upon priming in the presence of m+4Met and dC[D$_8$].

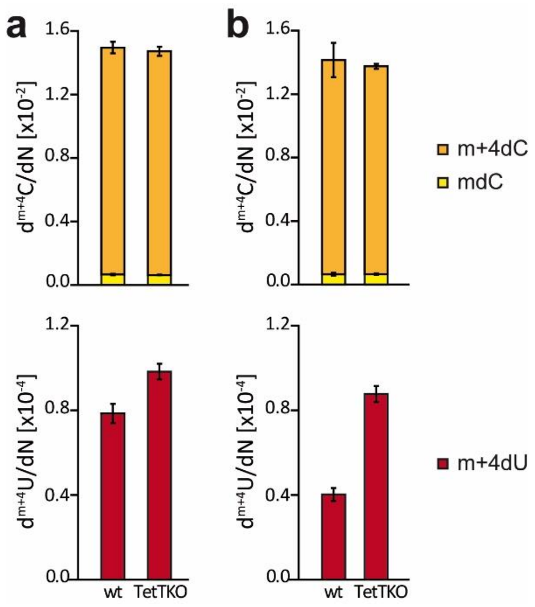

**Figure S7. Tet proteins do not trigger DNA repair events that contribute to turnover of mdC into dT** (related to Fig. 3)**.**

Global levels of mdC, m+4dC and m+4dU (red) in the genome of paretal (Tet1-3 proficient) and Tet TKO E14tg2a mPSCs after priming for five days in the presence m+4Met. **a**) and **b**) show indipendent biological replicates. Mean and standard deviation of three technical replicates are shown.

#### Supplementary Table 1

Oligonucleotide/primer sequences used in this study. RT = Reverse Transcription.

| Name | Sequence | Application | Reference |
|---|---|---|---|
| mCda gRNA F | CACCGAGCAGCCGCTGGACGTGCTC | gRNA construct | this work |
| mCda gRNA R | AAACGAGCACGTCCAGCGGCTGCTC | gRNA construct | this work |
| mDctd gRNA F | CACCGACATTCGTTACACGGGAACA | gRNA construct | this work |
| mDctd gRNA R | AAACTGTTCCCGTGTAACGAATGTC | gRNA construct | this work |
| mCda F | AAGGCCATCTCCGAAGGGTA | RT-PCR | this work |
| mCda R | TCTTCAGGTCCAAACGAGGC | RT-PCR | this work |
| Dnmt3b Fw | CTGGCACCCTCTTCTTCATT | RT-qPCR | 1 |
| Dnmt3b Rv | ATCCATAGTGCCTTGGGACC | RT-qPCR | 1 |
| Fgf5 Fw | AGGGGATTGTAGGAATACGAG | RT-qPCR | 3 |
| Fgf5 Rv | TCTTGGAATCTCTCCCTGAAC | RT-qPCR | 3 |
| Klf4 Fw | CGGGAAGGGAGAAGACACT | RT-qPCR | 3 |
| Klf4 Rv | GAGTTCCTCACGCCAACG | RT-qPCR | 3 |
| Nanog Fw | AAGTACCTCAGCCTCCAGCA | RT-qPCR | 4 |
| Nanog Rv | GCTTGCACTTCATCCTTTGG | RT-qPCR | 4 |
| Otx2 Fw | CATGATGTCTTATCTAAAGCAACCG | RT-qPCR | 5 |
| Otx2 Rv | GTCGAGCTGTGCCCTAGTA | RT-qPCR | 5 |
| Oct6 Fw | TTTCTCAAGTGTCCCAAGCC | RT-qPCR | 3 |
| Oct6 Rv | ACCACCTCCTTCTCCAGTTG | RT-qPCR | 3 |
| Oct4 Fw | AGGAAGCCGACAACAATGAG | RT-qPCR | 4 |
| Oct4 Rv | ACTCCACCTCACACGGTTCT | RT-qPCR | 4 |
| Prdm14 Fw | CTCTGGAGACAGGCCATACC | RT-qPCR | 3 |
| Prdm14 Rv | GTCTGATGTGTGTTCGGAGTATGC | RT-qPCR | 3 |
| Tfcp2l1 Fw | GGGGACTACTCGGAGCATCT | RT-qPCR | 3 |
| Tfcp2l1 Rv | TTCCGATCAGCTCCCTTG | RT-qPCR | 3 |
| Rex1 Fw | GCTCCTGCACACAGAAGAAA | RT-qPCR | 5 |
| Rex1 Rv | GTCTTAGCTGCTTCCTTCTTGA | RT-qPCR | 5 |
| Stella Fw | AAAGTCGACCCAATGAAGGA | RT-qPCR | 4 |
| Stella Rv | ACACCGGGGTTTAGGGTTAG | RT-qPCR | 4 |
| Gapdh Fw | TGGAGAAACCTGCCAAGTATGA | RT-qPCR | 1 |
| Gapdh Rv | GTTGCTGTTGAAGTCGCAGG | RT-qPCR | 1 |
| Dlk1 S1 F | GTCTCGTGGGCTCGGAGATGTGTATAAGAGACAG GTGTTTATTAGTTTGGTGGTG | Bsulfite PCR | this work |
| Dlk1 S1 R | CGACGCTCTTCCGATCTACTTAAATCTCCTCATCACCA | Bsulfite PCR | this work |
| H19 S2 F | GTCTCGTGGGCTCGGAGATGTGTATAAGAGACAG AAGAGAGAAGAAGGAGATTTATTTGT | Bsulfite PCR | this work |
| H19 S2 R | CGACGCTCTTCCGATCTCATCTAAAATCTAACAAAAATATTAAAAA | Bsulfite PCR | this work |
| H19 S3 F | GTCTCGTGGGCTCGGAGATGTGTATAAGAGACAG ATTATAATGGGAATTTGAGGGTA | Bsulfite PCR | this work |
| H19 S3 R | CGACGCTCTTCCGATCTTAACCAAAACCAAACTATAAAATAACTAAT | Bsulfite PCR | this work |
| Nesp S1a F | GTCTCGTGGGCTCGGAGATGTGTATAAGAGACAG TGAGTTTTTTGAATTTGAGTTTGAT | Bsulfite PCR | this work |
| Nesp S1a R | CGACGCTCTTCCGATCTTAAATAACTAATTAAAAATAACACCC | Bsulfite PCR | this work |
| Nesp S2 F | GTCTCGTGGGCTCGGAGATGTGTATAAGAGACAG GGAGGTTAAAGGTTTGTTGG | Bsulfite PCR | this work |
| Nesp S2 R | CGACGCTCTTCCGATCTACATAAAAAACAAATAACTTACCTTT | Bsulfite PCR | this work |
| MM S1b F | GTCTCGTGGGCTCGGAGATGTGTATAAGAGACAG TAGGGGATGGAGAAGGGT | Bsulfite PCR | this work |
| MM S1b R | CGACGCTCTTCCGATCTATTCTCCAACTTCTACTCTCA | Bsulfite PCR | this work |
| Cdkn1c 2 F | GTCTCGTGGGCTCGGAGATGTGTATAAGAGACAG GTTGTGAAATTGAAAATATTATATTATGTT | Bsulfite PCR | this work |
| Cdkn1c 2 R | CGACGCTCTTCCGATCTTAAATAAAACCCCTTACACAACC | Bsulfite PCR | this work |
| Nextera i7 | CAAGCAGAAGACGGCATACGAGATNNNNNNNNGTCTCGTGGGCTCGG | Indexing | this work |
| TruSeq i5 | AATGATACGGCGACCACCGAGATCTACACNNNNNNNN ACACTCTTTCCCTACACGACGCTCTTCCGATCT | Indexing | this work |

### SUPPLEMENTARY METHODS

#### High depth bisulfite amplicon sequencing

Genomic DNA (1-2 µg) was bisulfite treated using the EpiTect Bisulfite Kit (Qiagen). 5' overhangs compatible with Illumina TruSeq and Nextera adaptors were appended to the locus specific primers (Supplementary Table 1). Approximately 10 ng of bisulfite converted DNA were amplified with HotStar-Taq plus DAN polymerase (Qiagen). PCR products were analyzed by agarose gel electrophoresis and quantified by densitometry using ImageJ. Similar amounts of these primary amplicons were used as templates for a second round of amplification with i5 and i7 Indexing Primers (Supplementary Table 1). PCR products were again analyzed by garose gel electrophoresis and, after purification with CleanNGS magnetic beads (CleanNA), they were quantified by fluorometry with Quant-iT PicoGreen Reagent (Thermo Fisher). Equimolar amounts of amplicons were then pooled and the size distribution of the final library was assessed on a Bioanalyzer (Agilent). The 2x300 output mode for dual indexed sequencing was used on a MiSeq instrument (Illumina).

#### Reverse transcription (RT)-qPCR analysis

Total RNA was isolated from the lysate flow-throughs obtained upon genomic DNA isolation with silica spin columns. RNA isolation from the flow-throughs was performed as described in the Quick-DNA/RNA Miniprep kit (Zymo Research, D7001). In-column DNase I digestion was performed to minimize carryover of residual genomic DNA. One microgram of total RNA was reverse transcribed with the iScript cDNA Synthesis Kit and the cDNA product equivalent to 50 ng of total RNA were amplified using iTaq Univesal SYBR Green Supermix (both from BioRad) on a qTOWER$^3$/G cycler (Jena Biosciences). qPCR primers are reported in Supplementary Table 1. Each primer pair spanned across one intron except from Oct6 cDNA amplifications as Pou3f1 (the gene coding Oct6) is intronless. qPCR reactions were also performed in parallel for every target/sample by omitting the reverse transcriptase to control for signals generated from residual genomic DNA (intronless Oct6 transcript, relatively short introns and spliced pseudogenes). An RT-qPCR assay for Gapdh transcripts was used as house keeping transcript reference to calculate ΔCt values and expression fold change values thereof.

#### Chemical Synthesis

Unless noted otherwise, all reactions were performed using oven dried glassware under an atmosphere of argon. Molsieve-dried solvents were used from *Sigma Aldrich* and chemicals were bought from *Sigma Aldrich*, *TCI*, *Carbolution* and *Carbosynth*. For extraction and chromatography purposes, technical grade solvents were distilled prior to their usage. Reaction controls were performed using TLC-Plates from *Merck* (Merck 60 $F_{254}$), flash column chromatography purifications were performed on *Merck* Geduran Si 60 (40-63 µM). Visualization of the TLC plates was achieved through UV-absorption. NMR spectra were recorded in deuterated solvents on *Varian VXR400S*, *Varian Inova 400*, *Bruker AMX 600*, *Bruker Ascend 400* and *Bruker Avance III HD*. HR-ESI-MS spectra were obtained from a *Thermo Finnigan* LTQ FT-ICR. IR-measurements were performed on a *Perkin Elmer Spectrum BX*. HPLC purifications were performed on a *Waters Breeze* system (2487 dual array detector, 1525 binary HPLC pump) using a Nucleosil VP 250/10 C18 column from *Macherey Nagel*. HPLC-grade MeCN was purchased from *VWR*.

*General numbering of the pyrimidine nucleosides*

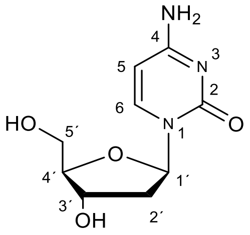

**Synthesis of 5-[$D_3$]-Methyl-2'-deoxy-[6-D, 1,3-$^{15}N_2$]-cytidine**

**[$D_2$]-Propionic acid**

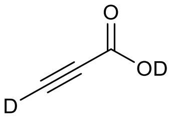

$Na_2CO_3$ (6.57 g, 62.0 mmol, 1.5 Equiv.) was dissolved in $D_2O$ (25 mL), propionic acid (2.60 mL, 42.0 mmol, 1.0 Equiv.) was added dropwise over 3 min. The reaction was stirred for 18 h at rt, the solvent was removed under reduced pressure, another portion of $D_2O$ (21 mL) was added and the mixture was stirred for 14 h at rt. Through addition of $D_2SO_4$ the pH value of the mixture was adjusted to 1. The mixture was extracted with DCM, the organic layers were dried over $MgSO_4$, all the volatiles were removed under reduced pressure (600 mbar, 40 °C). Purification by fractional distillation (27 mbar,39 °C) yielded [$D_2$]-propionic acid (452 mg, 6.27 mmol, 15 %) as a colorless liquid and was directly used for further reactions.

**[$D_2$,$^{15}N_2$]-Uracil**

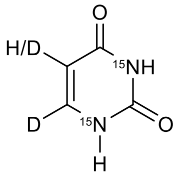

[$^{15}N_2$]-Urea (150 mg, 2.41 mmol, 1.0 Equiv.) was dissolved in $D_2SO_4$ (3 mL) and [$D_2$]-propiolic acid (0.161 mL, 2.65 mmol, 1.1 Equiv.) was added dropwise. The mixture was stirred at 105 °C for 18 h. The brown suspension was diluted with water (50 mL) and the pH value was adjusted to 10 with $Na_2CO_3$. Volatiles were removed under reduced pressure and the crude was purified by flash column chromatography (DCM/MeOH 4:1). A mixture of [$D_2$, $^{15}N_2$]-uracil (101 mg, 0.864 mmol, 36 %) und [D, $^{15}N_2$]-uracil (50 mg, 0.43 mmol, 18 %) was obtained as a colorless solid.

*$^1$H-NMR* (400 MHz, DMSO-d6) δ 11.03 (dd, $^1J$ = 77.2 Hz, $^4J$ = 1.8 Hz, 1H, N3-H), 10.80 (dd,$^1J$ = 82.6 Hz, $^4J$ = 1.9 Hz, 1H, N1-H), 5,46 (dd, $^3J$ = 4.3 Hz, $^4J$ = 2.5 Hz, C5-H) ppm.

*$^{13}$C-NMR* (101 MHz, DMSO-d6) δ 164.9 (t, $^1J$ = 9.7 Hz, C4), 151.9 (m, C2), 142.3 (m, C6), 101.4-99.5 (m, C5) ppm.

**[$D_2$,$^{15}N_2$]-3’,5’-Bis-O-(p-Toluoyl)-2’-deoxyuridine**

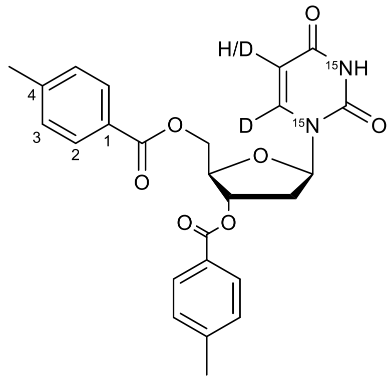

The mixture of [$D_2$, $^{15}N_2$]-uracil und [D, $^{15}N_2$]-uracil (37 mg, 0.32 mmol, 1.0 Equiv.) was dissolved in MeCN (2 mL). Bis-(trimethylsilyl)acetamide (0.3 mL, 0.96 mmol, 3.0 Equiv.) was added dropwise and the reaction mixture was stirred for 3 h at 80 °C. All volatiles were removed under reduced pressure and the residue was coevaporated with chloroform (3 x 6 mL). The residue was dissolved in chloroform(2 mL) and Hoffer’s chlorosugar (149 mg,0.383 mmol, 1.2 Equiv.) was added and stirred at rt for 17 h. After removing of the volatiles under reduced pressure, purification by flash column chromatography (*i*Hex/EtOAc 1:1) yielded a mixture of α-[$D_2$, $^{15}N_2$]-3’,5’-bis-O-(p-toluoyl)-2’-deoxyuridine β-[D, $^{15}N_2$]-3’,5’-bis-O-(p-toluoyl)-2’-deoxyuridine (41 mg, 0.088 mmol, 28 %, α/β: 1.28:1) as a yellow foam.

α-[$D_2$, $^{15}N_2$]-3’,5’-Bis-O-(p-toluoyl)-2’-deoxyuridine:

*$^{1}$H-NMR* (400 MHz, CDCl3): δ = 9.06 (d, $^{1}J$ = 91.0 Hz, 1H, NH), 7.89 (d, $^{3}J$ = 8.0 Hz, 2H, Tol2), 7.78 (d, $^{3}J$ = 8.0 Hz, 2H, Tol2), 7.31-7.20 (m, 4H, Tol3), 6.31 (d, $^{3}J$ = 6.8 Hz, 1H, C1’- H), 5.72 (dt, $^{3}J$ = 4.7 Hz, $^{3}J$ = 2.4 Hz, 1H, C5-H), 5.61 (m, 1H, C3’-H), 4.86 (t, $^{3}J$ = 4.3 Hz, 1H, C4’-H), 4.54 (m, 2H, C5’-H), 3.01-2.89 (m, 1H, C2’-H), 2.55 (d, $^{3}J$ = 15.5 Hz, 1H, C2’-H), 2.43 (s, 3H, $CH_3$), 2.41 (s, 3H, $CH_3$) ppm.

*$^{13}$C-NMR* (101 MHz, CDCl3): δ = 165.7 (COO), 166.1 (COO), 150.1 (C2), 144.9 (Tol4), 144.4 (Tol4), 129.7 (Tol2), 129.6 (Tol2), 129.5 (Tol3), 129.4 (Tol3), 126.5 (Tol1), 125.9 (Tol1), 101.4 (C5), 88.0 (d, 1J = 10.0 Hz, C1’), 85.7 (C4’), 74.7 (C3’), 64.0 (C5’), 39.0 (C2’), 21.7 ($CH_3$) ppm.

β-[$D_2$, $^{15}N_2$]-3’,5’-Bis-O-(p-toluoyl)-2’-deoxyuridine:

*$^{1}$H-NMR* (400 MHz, CDCl3): δ = 8.93 (d, $^{1}J$ = 91.0 Hz, 1H, NH), 7.94 (d, $^{3}J$ = 8.0 Hz, 2H, Tol2), 7.93 (d, $^{3}J$ = 8.0 Hz, 2H, Tol2) 7.31-7.20 (m, 4H, Tol3), 6.41 (dd, $^{3}J$ = 8.3 Hz, $^{3}J$ = 5.7 Hz, 1H, C1’-H), 5.60 (m, 2H, C3’-H und C5-H), 4.73 (dd, $^{3}J$ = 12.3 Hz, $^{4}J$ = 3.1 Hz, 1H, C5’-H), 4.67 (dd, $^{3}J$ = 12.3 Hz, $^{4}J$ = 3.5 Hz, 1H, C5’-H), 4.60-4.49 (m, 1H, C4’-H), 2.75 (dd, $^{3}J$ = 14.3 Hz, $^{3}J$ = 5.7 Hz, 1H, C2’-H), 2.43 (s, 6H, CH3), 2.30 (m, 1H, C2’-H) ppm.

*$^{13}$C-NMR* (101 MHz, CDCl3): δ = 165.7 (COO), 166.1 (COO), 150.1 (C2), 144.7 (Tol4), 129.9 (Tol2), 129.5 (Tol2), 129.4 (Tol3), 129.3 (Tol3), 126.5 (Tol1), 125.9 (Tol1), 101.4 (C5), 85.4 (d, 1J = 12.8 Hz, C1’) 83.0 (C4’), 74.7 (C3’), 64.0 (C5’), 38.4 (C2’), 21.7 ($CH_3$) ppm.

*LRMS (ESI$^+$):* m/z calculated $[C_{25}H_{23}D_2O_7{}^{15}N_2]^+$ ($[M+H]^+$): 469.17, found: 469.15.

**β-[D, $^{15}N_2$]-3',5'-Bis-O-(p-toluoyl)-5-iodo-2'-deoxyuridine**

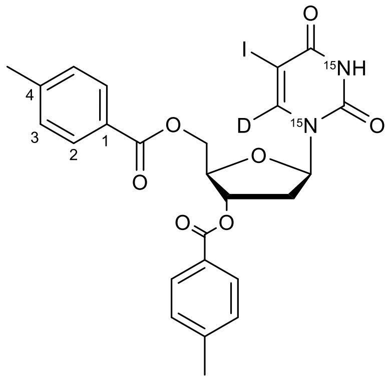

A mixture of α-[$D_2$, $^{15}N_2$]-3',5'-bis-O-(p-toluoyl)-2'-deoxyuridine and β-[$D_2$, $^{15}N_2$]-3',5'-bis-O-(p-toluoyl)-2'-deoxyuridine (267 mg, 0.570 mmol, 1.0 Equiv.) was dissolved in MeCN (6 mL), LiI (92 mg, 0.68 mmol, 1.2 Equiv.) and ceric(IV)-ammoniumnitrate (625 mg, 1.14 mmol, 2.0 Equiv.) were added. The suspension was stirred at 80 °C for 3 h. After full conversion of the starting material, saturated aq. NaCl-solution (30 mL) was added, mixture was extracted with DCM, the combined organic layers were dried oved $MgSO_4$ and all volatiles were removed under reduced pressure. Purification by flash column chromatography (*i*Hex/EtOAc 1:1) yielded β-[D, $^{15}N_2$]-3',5'-bis-O-(p-toluoyl)-5-iodo-2'-deoxyuridine 33 (177 mg,0.298 mmol, 52 %) as a yellow solid.
*$^1$H-NMR* (400 MHz, $CDCl_3$): δ = 8.99 (d, $^1J$ = 93.6 Hz, 1H, NH), 7.94 (d, $^3J$ = 3.5 Hz, 2H, Tol2), 7.92 (d, $^3J$ = 3.6 Hz, 2H, Tol2), 7.27 (d, $^3J$ = 7.7 Hz, 4H, Tol3), 6.37 (dd, $^3J$ = 8.7 Hz, $^3J$ = 5.4 Hz, 1H, C1'-H), 5.61 (d, $^3J$ = 6.6 Hz, 1H, C3'-H), 4.73 (t, $^3J$ = 2.7 Hz, 2H, C5'-H), 4.57 (q, $^3J$ = 2.7 Hz, 1H, C4'-H), 2.78 (dd, $^3J$ = 14.3, 3J = 5.5 Hz, 1H, C2'-H), 2.43 (s, 3H, $CH_3$), 2.42 (s, 3H, CH3), 2.28 (m, 1H, C2'-H) ppm.
*$^{13}$C-NMR* (101 MHz, $CDCl_3$): δ = 166.1 (COO), 166.0 (COO), 159.6 (d, $^1$J = 10.4 Hz, C4), 149.7 (t, $^1$J = 18.9 Hz, C2), 144.7 (Tol4), 144.6 (Tol4), 129.9 (Tol2), 129.7 (Tol2), 129.6 (Tol3), 129.3 (Tol3), 126.4 (Tol1), 126.2 (Tol1), 85.8 (d, $^1J$ = 12.3 Hz, C1'), 83.5 (C4'), 74.9 (C3'), 68.8 (d, $^2J$ = 9.0 Hz, C5), 64.2 (C5'), 38.8 (C2'), 21.8 ($CH_3$) ppm.
HRMS (ESI$^+$): m/z calculated $[C_{25}H_{23}DO_7{}^{15}N_2I]^+$ ($[M+H]^+$): 592.0480, found:592.0476.

**β-[D, $^{15}N_2$]-3',5'-Bis-O-(p-Toluoyl)-5-iodo-2'-deoxycytidine**

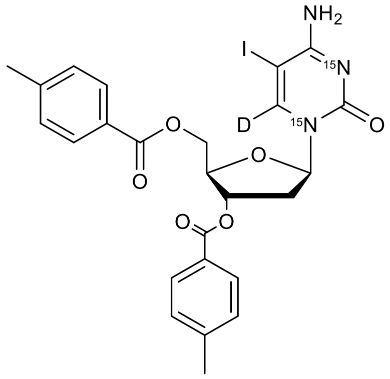

1,2,4-Triazole (42 mg, 0.60 mmol, 9.0 Equiv.) was dissolved in MeCN (1 mL), $POCl_3$ (0.028 mL, 0.29 mmol, 4.4 Equiv.) was added dropwise at 0 °C, the mixture was stirred for 10 min at 0 °C, $NEt_3$ (0.084 mL,0.60 mmol, 9.0 Equiv.) was added and the reaction solution was stirred at rt for 20 min. A solution of β-[D, $^{15}$N2,]-3',5'-bis-O-(p-Toluoyl)-5-iodo-2'-deoxyuridine (40 mg, 0.07 mmol, 1.0 Equiv.) in MeCN (1.5 mL) was added to the reaction and stirred at 30 °C for 18 h. After full conversion of the starting material, $NEt_3$ (0.4 mL) und $H_2O$ (0.1 mL) were added and stirred for 10 min at 30 °C. Saturated aq. NaCl-solution (5 mL) and

saturated aq. $NaHCO_3$-solution (5 mL) were cooled down to 0 °C and added to the reaction mixture. The solution was extracted with DCM, the combined organic layers were dried over $MgSO_4$ and all volatiles were removed under reduced pressure. The residue was dissolved in 1,4-dioxane (10 mL) and a concentrated $NH_3$-solution (1.5 mL) was added. The reaction mixture was stirred for 15 min at 30 °C, afterwards an aqueous $NH_4Cl$-solution (10 mL) was added and extracted with DCM. The combined organic layers were dried over $MgSO_4$ and the volatiles were removed under reduced pressure. Purification of the crude by flash column chromatography (DCM/MeOH 50:1) yielded [D, $^{15}N_2$]-3',5'-bis-O-(p-toluoyl)-5-iodo-2'-deoxycytidine (17 mg, 0.029 mmol, 43 %) as a yellow solid.
*$^{1}H$-NMR* (400 MHz, $CDCl_3$): δ = 7.94 (d, $^{3}J$ = 8.0 Hz, 2H, Tol2), 7.90 (d, $^{3}J$ = 8.1 Hz, 2H, Tol2), 7.29-7.22 (m, 4H, Tol3), 6.36 (dd, $^{3}J$ = 8.4 Hz, $^{3}J$ = 5.4 Hz, 1H, C1'-H), 5.58 (d, $^{3}J$ = 6.4 Hz, 1H, C3'-H), 5.49 (s, 2H, NH2), 4.76 (dd, $^{3}J$ = 12.3 Hz, $^{3}J$ = 3.0 Hz, 1H, C5'-H), 4.68 (dd, $^{3}J$ = 12.3 Hz, $^{3}J$ = 3.4 Hz, 1H, C5'-H), 4.59 (q, $^{3}J$ = 2.7 Hz, 1H, C4'-H), 3.02-2.92 (m, 1H, C2'-H), 2.42 (s, 3H, $CH_3$), 2.41 (s, 3H, $CH_3$), 2.17 (m, 1H, C2'-H) ppm.
*$^{13}C$-NMR* (101 MHz, CDCl3): δ = 166.1 (COO), 163.7 (C4), 154.9 (C2), 144.5 (Tol4) 144.5 (Tol4), 129.8 (Tol2), 129.7 (Tol2), 129.5 (Tol3), 129.3 (Tol3), 126.4 (Tol1), 126.3 (Tol1), 87.1 (d, $^{1}J$ = 11.2 Hz, C1'), 83.6 (C4'), 75.3 (C3'), 64.2 (C5'), 56.1 (C5), 39.5 (C2'), 21.8 ($CH_3$) ppm.
HRMS (ESI$^+$): m/z calculated $[C_{25}H_{24}DO_6N^{15}N_2I]^+$ ($[M+H]^+$): 593.0792, found: 593.0791.

**5-[$D_3$]-Methyl-2'-deoxy-[6-D, 1,3-$^{15}N_2$]-cytidine**

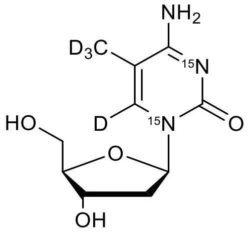

$Ni(dppp)Cl_2$ (8 mg, 0.01 mmol, 0.5 Equiv.) was dried in a schlenk flask under high vacuum for 30 min. A solution of β-[D, $^{15}N_2$]-3',5'-bis-O-(p-toluoyl)-5-iodo-2'-deoxycytidine (17 mg, 0.029 mmol, 1.0 Equiv.) in THF (1 mL) was added at 0 °C. Afterwards $D_3CMgI$ (0.072 mL, 0.072 mmol, 2.5 Equiv.) was added dropwise and the mixture was stirred at rt for 1.5 h. Another portion of $D_3CMgI$ (0.072 mL, 0.072 mmol, 2.5 Equiv.) was added dropwise and stirred for 1 h at rt. A third portion of $D_3CMgI$ (0.20 mL, 0.20 mmol, 6.9 Equiv.) and $Ni(dppp)Cl_2$ (8 mg, 0.01 mmol, 0.5 Equiv.) were added to the mixture and stirred for17 h at rt. A saturated aqueous $NH_4Cl$-solution (8 mL) was added to the reaction solution and extracted with EtOAc. The combined organic layers were dried $MgSO_4$ and all volatiles were removed under reduced pressure The residue was purified by flash column chromatography (DCM/MeOH 20:1) to yield a mixture of β-5-[$D_3$]-methyl-3',5'-bis-O-(p-toluoyl)-2'-deoxy-[D,$^{15}N_2$,]-cytidine and β-3',5'-bis-O-(p-toluoyl)-2'-deoxy-[D, $^{15}N_2$,]-cytidine as a yellow foam.
The resulting mixture was dissolved in MeOH (1.2 mL), $K_2CO_3$ (13 mg, 0.094 mmol, 4.7 Equiv.) was added and the solution was stirred for 18 h at rt. Another portion of $K_2CO_3$ (13 mg, 0.094 mmol, 4.7 Equiv.) and MeOH (1.2 mL) was added and stirred for 17 h at rt. The volatiles were removed under reduced pressure and the residue was purified via HPLC (Macherey-Nagel, Nucleosil 100-7 C18, 10 × 250 mm, linear gradient, 0 % - 10 % MeCN in water in 45 min) to yield [$D_4$, 15$N_2$]-m5dC (0,9 mg,0,004 mmol, 12 %) as a colorless solid.
*$^{1}H$-NMR* (800 MHz, $D_2O$): δ = 6.30 (t, $^{3}J$ = 6.7 Hz, 1H, C1'-H), 4.46 (dt, $^{3}J$ = 6.6 Hz, $^{3}J$ = 4.1 Hz, 1H, C3'-H), 4.05 (q, $^{3}J$ = 4.2 Hz, 1H, C4'-H), 3.86 (dd, $^{3}J$ = 12.5 Hz, $^{3}J$ = 3.6 Hz, 1H, C5'-H), 3.78 (dd, $^{3}J$ = 12.5 Hz, $^{3}J$ = 5.1 Hz, 1H, C5'-H), 2.47-2.36 (m, 1H, C2'-H), 2.38-2.25 (m, 1H, C2'-H) ppm.
*$^{13}C$-NMR* (201 MHz, $D_2O$): δ = 89.1 (C4'), 88.4 (d, $^{1}J$ = 13.0 Hz, C1'), 73.1 (C3'), 63.8 (C5'), 41.8 (C2') ppm.
HRMS (ESI$^+$): m/z calculated $[C_{10}H_{12}D_4O_4N^{15}N_2]$+ ([M+H]+): 248.1327, found: 248.1326.

**Synthesis of 5-formyl-2'-deoxy-(1',2',3',4',5'-$^{13}C_5$, *N$^1$,N$^3$*-$^{15}N_2$)-cytidine**

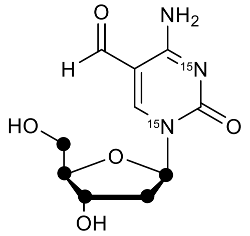

5-formyl-2'-deoxy-(1',2',3',4',5'-$^{13}C_5$, N1,N3-$^{15}N_2$)-cytidine was prepared according to Iwan *et al.*[6]

##### 1.3 Synthesis of 5-hydroxymethyl-2'-deoxy-(1',2',3',4',5'-$^{13}C_5$, *N$^1$,N$^3$*-$^{15}N_2$)-cytidine

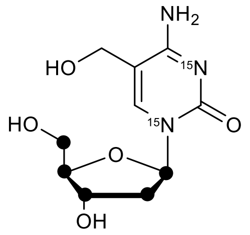

5-formyl-2'-deoxy-(1',2',3',4',5'-$^{13}C_5$, N1,N3-$^{15}N_2$)-cytidine (7.4 mg, 0.028 mmol, 1.0 Equiv.), $CeCl_3$*$7H_2O$ (32 mg, 0.085 mmol, 3.0 Equiv.) and $NaBH_4$ (1.0 mg, 0.028 mmol, 1.0 Equiv.) were dissolved in MeOH and stirred for 5 min at rt. Saturated aqeous $NH_4Cl$ solution was added and the mixture was concentrated under reduced pressure. Purification via HPLC (Macherey-Nagel, Nucleosil 100-7 C18, 10 × 250 mm, linear gradient, 0 % - 10 % MeCN in water in 60 min) yielded 5-hydoxymethyl-2'-deoxy-(1',2',3',4',5'-$^{13}C_5$, N1,N3-$^{15}N_2$)-cytidine (quant.).
HRMS (ESI$^+$): m/z calculated $[C_5{}^{13}C_5H_{16}O_5N^{15}N_2]^+$ ($[M+H]^+$): 265.1193, found: 265.1190.

#### 1.4 Synthesis of 5-methyl-2’-deoxy-(1’,2’,3’,4’,5’-$^{13}C_5$, *N$^1$,N$^3$*-$^{15}N_2$)-uridine

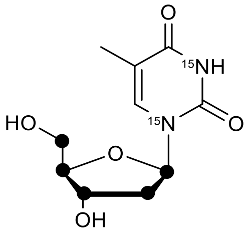

The detailed synthesis of this compound will be published in a forthcoming manuscript.
